# Stanniocalcin 1a is a Ca^2+^-regulated switch controlling epithelial cell quiescence-proliferation balance and Ca^2+^ uptake

**DOI:** 10.1101/2020.09.09.290114

**Authors:** Shuang Li, Chengdong Liu, Allison Goldstein, Yi Xin, Caihuan Ke, Cunming Duan

## Abstract

The mechanisms governing cell quiescence-proliferation balance are poorly defined. Using a zebrafish model, here we report that Stc1a, a glycoprotein known as a hypocalcemic hormone, not only inhibits epithelial calcium uptake but also functions as a Ca^2+^-regulated switch controlling epithelial cell quiescence-proliferation balance. Among the 4 *stc* genes, only the *stc1a* expression is [Ca^2+^]-dependent. Genetic deletion of *stc1a*, but not *stc2b*, resulted in elevated body Ca^2+^ contents, ectopic Ca^2+^ deposit, body swelling, and premature death. Reducing epithelial calcium channel Trpv6-mediated Ca^2+^ uptake alleviated these phenotypes. Loss of Stc1a also promoted quiescent epithelial cells to re-enter the cell cycle. This action was accompanied by local IGF signaling activation and increased expression in *papp-aa*, a zinc metalloproteinase degrading Igfbp5a. Genetic deletion of *papp-aa* or *igfbp5a* abolished the elevated epithelial cell reactivation in *stc1a^-/-^* mutants. Likewise, inhibition of IGF1 receptor, PI3 kinase, Akt, and Tor signaling abolished epithelial cell reactivation. These results reveal that Stc1a plays dual roles in regulating epithelial calcium uptake and cell quiescence-proliferation balance and implicate Trpv6 and Papp-aa-Igfbp5a-IGF signaling in these functions.

## Introduction

The majority of cells in the human body is in the non-proliferative quiescent state. Prominent examples include post-mitotic neurons and skeletal muscle cells. While these terminally cells can no longer re-enter the cell cycle, others, such as adult stem cells, hepatocytes, epithelial cells, and T cells, retain the capacity to re-enter the cell cycle in response to appropriate stimuli (Cheung and Rando, 2013). Maintaining a pool of quiescent cells that can be rapidly reactivated upon appropriate stimulation is critical for tissue repair, wound healing, and regeneration (Cheung and Rando, 2013). Dysregulation of the quiescence-proliferation balance are linked to human diseases such as cancer, autoimmune diseases, and fibrosis (Fiore et al., 2018; Kitaori et al., 2009). Historically, cellular quiescence is considered a passive outcome of cell cycle exit. Recent studies, however, suggest that cellular quiescence is highly regulated (Lacorazza, 2018). Studies in genetically tractable organisms have shown that the nutrient sensitive insulin/insulin-like growth factor (IGF)-PI3 kinase-AKT-mTOR signaling pathway plays a key role in this regulation. *Drosophila* neural stem cells are reactivated in response to dietary amino acids due to increased insulin release from neighboring glia cells (Britton and Edgar, 1998; Chell and Brand, 2010; Huang and Wang, 2018; Sousa-Nunes et al., 2011). Activation of mTOR signaling by deletion of the Tsc gene in adult mouse adult hematopoietic stem cells (HSCs) promotes these quiescent cells for cell cycle reentry and proliferate (Chen et al., 2008). Conversely, inhibition of mTOR activity preserved the long-term self-renewal and the hematopoietic capacity of HSCs (Chen et al., 2009). Mouse genetic studies revealed that IGF2 plays a key role in reactivating HSCs, neural stem cell, and intestinal stem cells (Ferron et al., 2015; Venkatraman et al., 2013; Ziegler et al., 2019; Ziegler et al., 2015). This regulation is not limited to adult stem cells. Naive T cells circulate in the body as quiescent cells with characteristically low mTOR activity. When presented with an antigen, mTORC1 signaling is activated in these cells, promoting them to exit quiescence and rapidly proliferate (Yang et al., 2013). While the importance of the insulin/IGF-PI3 kinase-AKT-mTOR signaling pathway in regulating the quiescence-proliferation balance is clear, how this central signaling pathway is activated in such a cell type- and physiological/pathological state-specific manner is poorly understood.

Our recent studies using a zebrafish model suggested a similar role of IGF-PI3 kinase-Akt-Tor signaling in regulating the proliferation-quiescence balance in a population of epithelial cells (Dai et al., 2014; Liu et al., 2017). These mitochondria-rich cells, known as ionocytes or NaR cells, take up Ca^2+^ from the surrounding aquatic habitat to maintain body Ca^2+^ homeostasis (Hwang, 2009; Yan and Hwang, 2019). In the adult stage, NaR cells are distributed in the adult gills, intestine, and kidney, key organs for Ca^2+^ uptake and reabsorption. In embryos and larval stages, however, these cells are distributed in the yolk sac epidermis (Hwang, 2009), making them highly accessible to experimental manipulation and observation. When zebrafish larvae are kept in media containing physiological concentrations of Ca^2+^, NaR cells are largely quiescent and IGF signaling activity in these cells are very low. When subjected to low [Ca^2+^] stress, these cells rapidly re-enter the cell cycle and proliferate due to elevated IGF-PI3 kinase-Akt-Tor signaling (Dai et al., 2014; Liu et al., 2017). Further studies suggested that IGF binding protein 5a (Igfbp5a), a secreted high-affinity IGF binding protein, and its major proteinase, pregnancy-associated plasma protein-a (Papp-aa), are highly expressed in NaR cells and indispensable in the low [Ca^2+^] stress-induced activation of IGF signaling in these cells (Liu et al., 2020; Liu et al., 2018). Genetic deletion of *igfbp5a, papp-aa*, or perturbations of Papp-aa-mediated Igfbp5a proteolysis abolishes NaR cell reactivation (Liu et al., 2020; Liu et al., 2018). Interestingly, the Papp-aa-mediated Igfbp5a proteolytic cleavage is inhibited by an unknown post-transcriptional mechanism under normal [Ca^2+^] conditions. Under low [Ca^2+^] conditions, this suppression mechanism is removed, which liberates IGFs from the Igfbp5a/IGF complex to bind and activate the IGF1 receptor (Liu et al., 2020). The molecular nature of this [Ca^2+^]-dependent mechanism, however, is unknown.

Stanniocalcin 1 (Stc1) is a dimeric glycoprotein originally discovered in the corpuscle of Stannius (CS), a fish-specific organ first described by Hermann Stannius in 1839 (Wagner and Dimattia, 2006). Early studies showed that surgical removal of CS led to increased blood calcium level and elevated calcium uptake (Fenwick, 1974; Fontaine, 1964; Pang, 1971), leading to the notion that CS contains a hypocalcemic factor (Wagner and Dimattia, 2006). The active CS component was purified and named as Stc1 (Wagner and Dimattia, 2006; Yeung et al., 2012). Mature Stc1 contains 11 conserved cysteine residues and a N-linked glycosylation site. The first 10 cysteines form intramolecular disulfide bridges and the 11th cysteine forms disulfide bond linking the two monomers, which stabilizes the functional dimer (Yamashita et al., 1995). Although Stc1 was considered to be a teleost fish-specific hormone for several decades and were once named teleocalcin, it is now appreciated that that multiple *Stc/STC* genes are present in all vertebrates including humans (Gagliardi et al., 2005; Wagner and Dimattia, 2006; Yeung et al., 2012). Humans have two *STC* genes, *STC1* and *STC2*, while zebrafish have 4 *stc* genes (Schein et al., 2012). These zebrafish genes are referred by different names in the literature and databases. Hereafter, they will be referred as *stc1a, stc1b, stc2a*, and *stc2b* following Schein et al. (2012) (Schein et al., 2012). In zebrafish, forced expression and morpholino-based knockdown of Stc1a changed Ca^2+^ uptake and altered NaR cell number, although these manipulations also changed the number of other ionocyte types and the uptake of other ions (Chou et al., 2015; Tseng et al., 2009). There is limited and conflicting evidence on whether mammalian STC1 regulates calcium uptake (Gagliardi et al., 2005; Wagner and Dimattia, 2006; Yeung et al., 2012). In contrast, many studies have linked human STC1 and STC2 to IGF signaling (Argente et al., 2017). In vitro biochemical studies showing that STC1 and STC2 can bind to and inhibit PAPP-A’s proteinase activity in cleaving IGFBPs (Argente et al., 2017). Overexpressing human STC1 or STC2 in transgenic mice resulted in reduced body size (Johnston et al., 2010). While the Stc1 knockout mice did not show notable growth abnormalities (Johnston et al., 2010), Stc2 knockout mice increased body size (Chang et al., 2008). Clinical studies have shown that human individuals carrying loss-of-function mutations in the *STC2* gene had greater adult height and this was linked to reduced PAPP-A activity and increased local IGF signaling activity (Marouli et al., 2017). These observations have led us to hypothesize that one or more of the Stc protein functions as [Ca^2+^]-dependent regulator of NaR cell quiescence and it acts by regulating IGF signaling. The objective of this study is to test this hypothesis.

## Results

### The expression of *stc1a* is regulated by Ca^2+^ levels

To determine the effects of Ca^2+^ in regulating *stc* gene expression, zebrafish were raised in embryo media containing varying concentrations of [Ca^2+^] (Figure 1A). The levels of *stc1a* mRNA, but not those of *stc1b, stc2a*, and *stc2b*, were [Ca^2+^]-dependent (Figure 1A-1D). Similar results were obtained in larval fish (Figure 1-figure supplement 1A). Next, *stc* mRNA levels in *trpv6^-/-^* and *papp-aa^-/-^* mutant fish were determined. Both mutant lines suffer from severe body calcium deficiency (Liu et al., 2020; Xin et al., 2019). The levels of *stc1a* mRNA, but not the other *stc* genes, were significantly reduced in the *papp-aa^-/-^* mutant fish compared to the wildtype fish (Figure 1E-H). Similarly, *stc1a* mRNA level was significantly decreased in the *trpv6^-/-^* mutant fish (Figure 1-figure supplement1B). These data suggest that the expression of *stc1a*, but not other *stc* genes, is regulated in a [Ca^2+^]-dependent manner.

**Figure 1.**
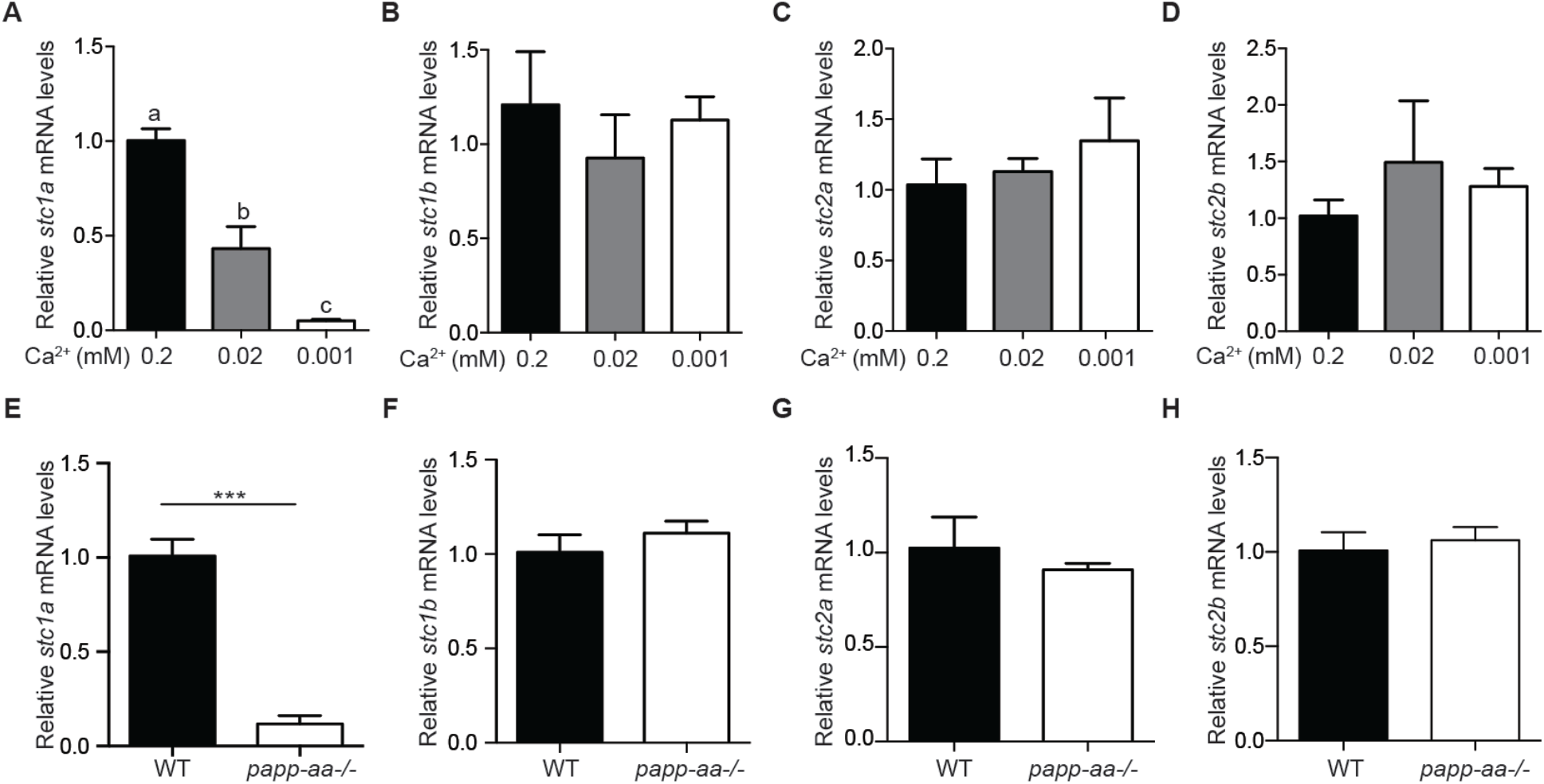
The expression of *stc1a*, but not other *stc* genes, is regulated by Ca^2+^ states. (**A-D**) Wild-type zebrafish embryos were raised in embryo media containing the indicated Ca^2+^ concentration until 3 days post fertilization (dpf). The mRNA expression levels of *stc1a* (**A**), *stc1b* (**B**), *stc2a* (**C**) and *stc2b* (**D**) were determined by qPCR and normalized by β-actin mRNA levels. Data shown are from 3 independent experiments, each containing 10-15 larvae/group. In this and all subsequent figures, data shown are Mean ± SEM unless stated otherwise. Different letters indicate significant differences between groups by one-way ANOVA followed by Tukey’s multiple comparison test (*P* < 0.05) unless stated otherwise. (**E-H)** Zebrafish embryos of the indicated genotypes were raised in E3 embryo medium. The mRNA levels of *stc1a* (**E**), *stc1b* (**F**), *stc2a* (**G**) and *stc2b* (**H**) were measured, normalized, and shown. Data shown are from 3 independent experiments, each containing 10-15 larvae/group. ***, *P* < 0.001 by unpaired twotailed t test.

### Genetic ablation of *stc1a* leads to cardiac edema, body swelling, and premature death

Three *stc1a* mutant fish lines, i.e., *stc1a (+17), stc1a (Δ18+1)*, and *stc1a (Δ35)*, were generated using CRISPR/Cas 9 (Figure 2- Figure 2-figure supplement 1). All 3 mutants lack the majority of the protein coding region and are predicted to be null mutations (Figure 2A). Progenies of *stc1a (+17)^+/-^* intercrosses, which were a mixture of homozygous, heterozygous, and wild type embryos, were raised and analyzed in a blind fashion at various stages followed by individual genotyping. No difference in the body size, development speed, and gross morphology was detected among the 3 genotypes until 3 dpf (Figure 2B-F). At 4 dpf, *stc1a (+17)^-/-^* mutant fish developed cardiac edema (Figure 2B). At 5 dpf, the cardiac edema became more pronounced. Additionally, mutant fish had no inflated swim bladder and displayed impaired touch-swim response (Figure 2B and Figure 2-figure supplement 2). By 6 and 7 dpf, mutant fish showed severe body swelling and death rate increased (Figure 2B and 2G). All mutant fish died by 10 days (Figure 2G). Similar results were obtained from the *stc1a (Δ18+1)^-/-^* fish line (Figure 2-figure supplement 3). In comparison, genetic deletion of *stc2b* did not cause morphological defects or premature death (Figure 2-figure supplement 4 and 5). These findings suggest that while Stc1a does not affect embryonic pattering and growth, it is essential for larval survival.

**Figure 2.**
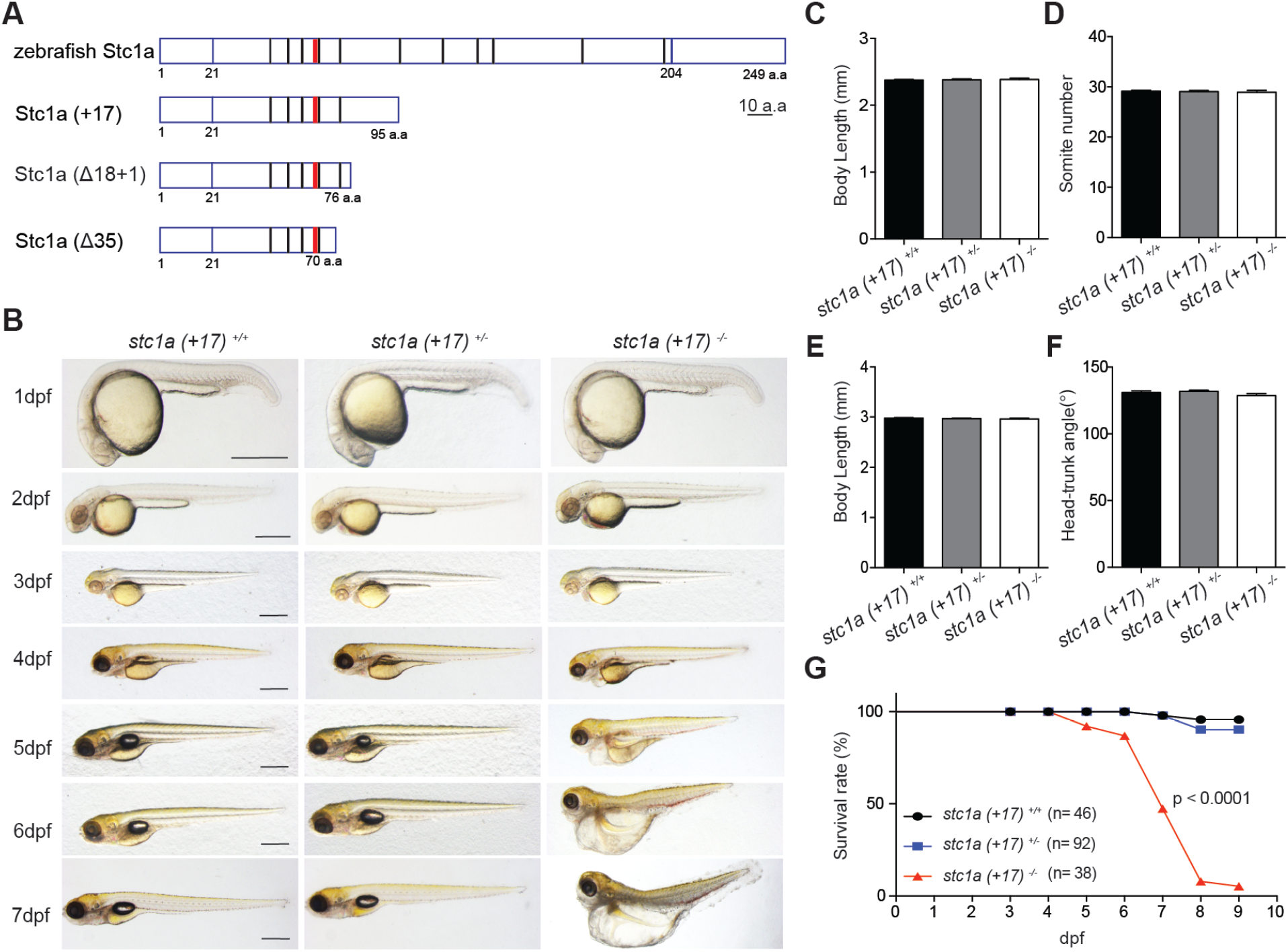
Genetic deletion of *stc1a* leads to cardiac edema, body swelling, and premature death. (**A**) Schematic diagram of Stc1a protein and the indicated mutants. The N-linked glycosylation site and conserved cysteine residues are shown by red and black bars, respectively. (**B)** Gross morphology of fish at the indicated stages. Lateral views with anterior to the left and dorsal up. Scale bar = 0.5 mm. (**C-F**) Body length, somite number, and head-trunk angle of the indicated genotypes at 24 hpf (**C, D**) and 48 hpf (**E, F**). n = 15-42 larvae/group. (**G**) Survival curves. Progenies of *stc1a (+17)^+/-^* intercrosses were raised in E3 embryo medium. Dead embryos were collected daily and genotyped individually. The survival curves of indicated genotypes and the total fish numbers are shown. *P* < 0.0001 by log-rank test.

### Loss of *stc1a* results in ectopic calcification and body edema by increasing Trpv6-mediated Ca^2+^ uptake

Because the previously reported role of Stc1 in regulating calcium homeostasis, we measured body Ca^2+^ levels in the mutant fish. Measuring blood Ca^2+^ levels in zebrafish embryos/larvae is technically not feasible due to their tiny size. Compared to their siblings, the total body Ca^2+^ levels in *stc1a^-/-^* mutants were significantly greater (Figure 3A). Alizarin red staining showed many strongly stained puncta in the yolk sac regions in the mutants at 5 and 7 dpf, indicating ectopic calcium deposits (Figure 3B). Furthermore, kidney stones like Ca^2+^ puncta were observed in pronephros in the mutant fish (Figure 3B). Similar data were obtained with the *stc1a (Δ18+1)^-/-^* fish (Figure 3-figure supplement 1A). In comparison, no ectopic Ca^2+^ puncta or kidney stone were observed in *stc2b^-/-^* mutant fish (Figure 3-figure supplement 1B).

**Figure 3.**
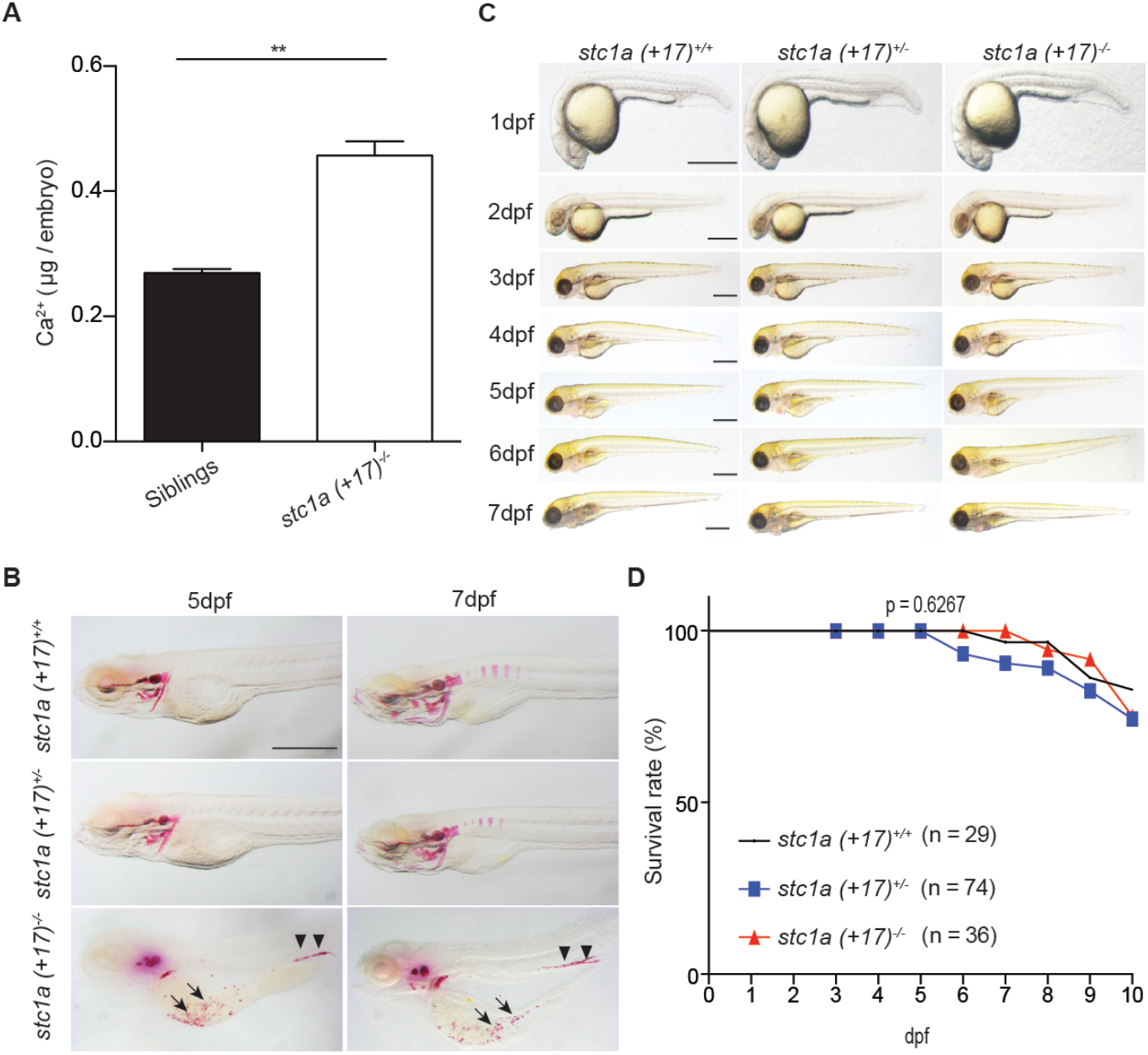
Abormal Ca^2+^ balance, cardiac edema, body swelling, and premature death in *stc1a^-/-^* fish are [Ca^2+^]-dependent. **(A)** Total body Ca^2+^ contents in 5 dpf zebrafish larvae of the indicated genotypes. Data shown are from 3 independent experiments, each containing 35 larvae/group. * *P*< 0.05 by unpaired two-tailed t test. (**B**) Zebrafish larvae of the indicated genotypes were raised in E3 embryo medium and subjected to Alizarin red staining at the indicated stages. Representative images are shown. Lateral view is shown with head to the left. Scale bar = 0.5 mm. Note the ectopic calcification in the yolk sac region (arrows) and kidney (arrow heads) in the mutant fish. (**C**) Gross morphology of fish at the indicated stages raised under low [Ca^2+^]. After raised in E3 embryo medium till 3 dpf, fish were transferred to the low [Ca^2+^] embryo medium and raised until the indicated stages. Representative morphological images of each genotype at the indicated stages are shown. Scale bar = 0.5 mm. (**D**) Survival curves under low [Ca^2+^]. After raised in E3 embryo medium till 3 dpf, they were transferred to the low [Ca^2+^] embryo medium from 3 dpf to 10 dpf. Dead fish were collected daily and genotyped individually. No statistical significance was detected among the three genotypes by log-rank test.

While the higher body Ca^2+^ contents, ectopic calcification, and kidney stones are consistent with the notion that Stc1a is a hypocalcemic hormone, the cardiac edema, body swelling, and premature death were unexpected. We postulated that the edema, body swelling, and premature death may be caused by the accumulation of osmotic water due to the increased osmotic influx and/or impaired kidney function. If this were correct, then removal of Ca^2+^ from the embryo rearing medium should alleviate the edema/swelling and premature death phenotypes. Indeed, Ca^2+^ depletion abolished the cardiac edema and body swelling in the mutant fish (Figure 3C). Ca^2+^ depletion also dramatically reduced ectopic calcification and kidney stone formation (Figure 3-figure supplement 2). Moreover, Ca^2+^ depletion treatment prevented the premature death caused by the loss of Stc1a, although it caused modest increases in all three groups (Figure 3D, Figure 3-figure supplement 3).

The epithelial calcium channel Trpv6 is specifically expressed in NaR cells and plays a key role in Ca^2+^ uptake (Dai et al., 2014; Xin et al., 2019). Gene expression analysis showed that the *trvp6* mRNA levels were significantly elevated in the mutant fish (Figure 4A). To determine whether the ectopic calcification observed in *stc1a^-/-^* fish is related to Trpv6-mediated Ca^2+^ uptake, progenies of *stc1a (+17)^+/-^* intercrosses were treated with CdCl_2_, a known Trpv6 inhibitor. CdCl_2_ treatment markedly reduced ectopic calcification and kidney stone formation (Figure 4B, Figure 4-figure supplement 1). Similar results were observed from GdCl_3_ treatment, another Trpv6 inhibitor (Figure 4-figure supplement 2A). Furthermore, CdCl_2_ treatment abolished cardiac edema and body swelling in *stc1a (+17)^-/-^* fish (Figure 4C) and rescued *stc1a (+17)^-/-^* fish from premature death (Figure 4D). Similarly, GdCl_3_ treatment significantly reduced cardiac edema and body swelling in *stc1a (+17)^-/-^* fish (Figure 4-figure supplement 2B). These results suggest that the Trpv6-mediated Ca^2+^ uptake is required for the aberrant calcium balance, the development of edema and body swelling, and premature death in *stc1a^-/-^* fish.

**Figure 4.**
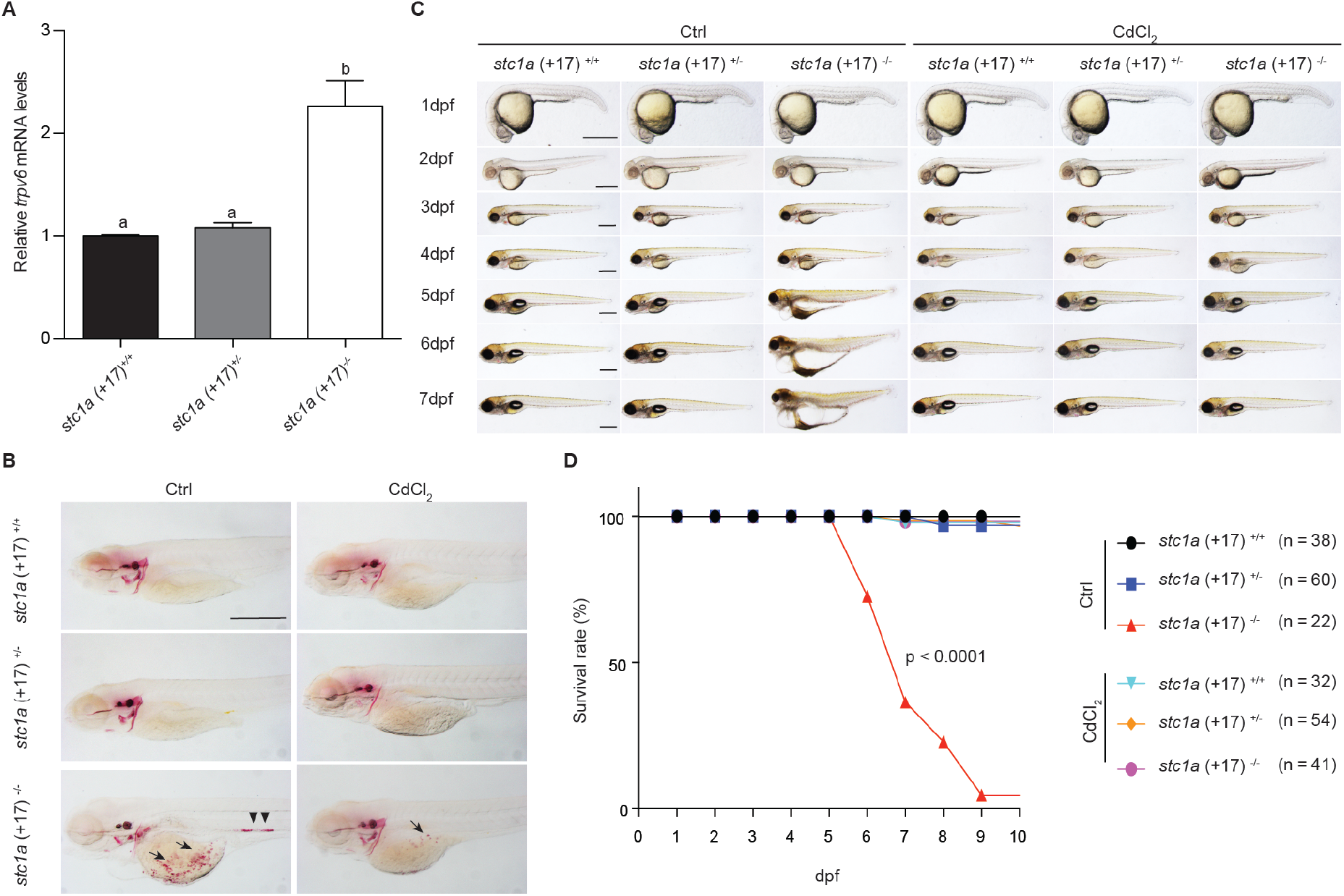
Stc1a regulates calcium balance and survival via Trpv6. (**A**) Increases *trpv6* mRNA levels in *stc1a^-/-^* mutants. Embryos of the indicated genotypes were raised in E3 embryo medium until 5 dpf. The mRNA levels of *trpv6* were measured and normalized. n = 15~17. **(B**) Progeny of *stc1a (+17)^+/-^* intercrosses were raised in E3 embryo medium and treated with or without 10 μg/L CdCl_2_ from 3 dpf to 5 dpf. The treated fish were subjected to Alizarin red staining. Fish was genotyped individually afterwards. Scale bar = 0.5 mm. Note the ectopic calcification in the yolk sac region (arrow) and kidney stones (arrow heads) in the mutant fish. **(C)** Progeny of *stc1a (+17)^+/-^* intercrosses were raised in E3 embryo medium and treated with or without 10 μg/L CdCl_2_ starting from 3 dpf until the indicated time. Representative morphological views of the indicated genotypes at the indicated stages are shown. Scale bar = 0.5 mm. **(D)** Inhibition of Trpv6 channel activity prevents *stc1a (+17)^-/-^* mutant fish from premature death. Progeny of *stc1a (+17)^+/-^* intercrosses were raised in E3 embryo medium and treated with or without 10 μg/L CdCl_2_ from 3 dpf until the indicated time. Dead fish were collected daily and genotyped individually. The survival curves of the indicated genotypes and the total fish numbers are shown. *P* < 0.0001 by log-rank test.

### Loss of Stc1a promotes NaR cell proliferation by activating IGF signaling

To test the hypothesis Stc1a functions as [Ca^2+^]-dependent regulator of NaR cell quiescence-proliferation balance, NaR cells in *stc1a (+17)^-/-^* mutants and siblings were labeled by in situ hybridization of *igfbp5a* mRNA, a specific NaR cell maker (Liu et al., 2017). *stc1a (+17)^-/-^* fish had significantly more NaR cells than their siblings (Figure 5A-B). Notably, NaR cells in the mutant fish were often observed as cell clusters, an indicator of newly divided cells (Figure 5A). The reactivation of NaR cells was confirmed by examining *stc1a (+17)^-/-^: Tg(igfbp5a:GFP)* and siblings (Figure 5D). *Tg(igfbp5a:GFP)* fish is a reporter fish line, in which NaR cells express EGFP (Liu et al., 2017). Similar elevation of NaR cell proliferation was observed in *stc1a (Δ18+1)* and *stc1a (Δ35)* mutant lines (Figure 5-figure supplement 1A-B). In contrast, no NaR cell reactivation was found in *stc2b^-/-^* fish (Figure 5-figure supplement 1C).

**Figure 5.**
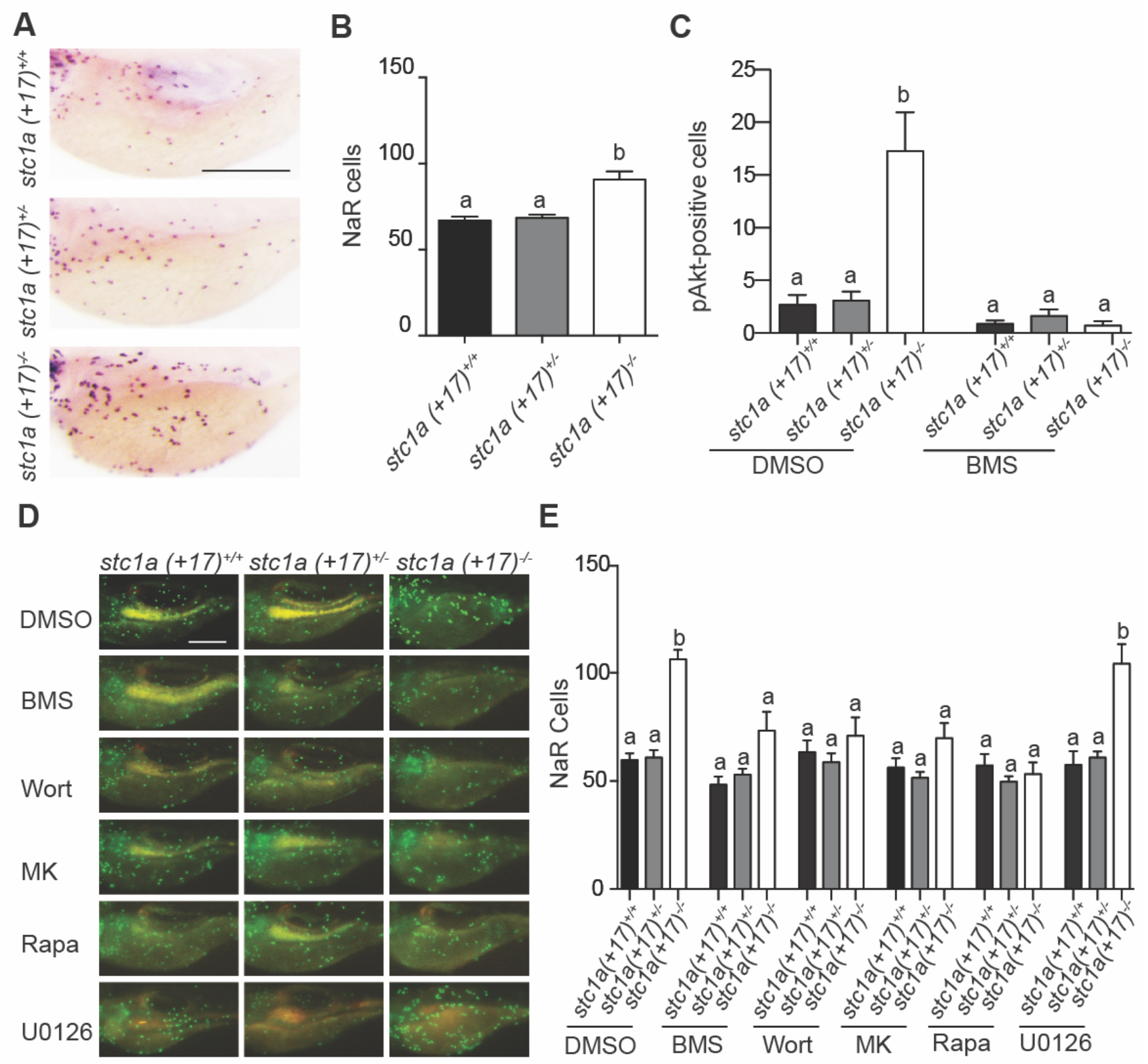
Genetic deletion of *stc1a* promotes NaR cell proliferation via regulating IGF-PI3 kinase-Akt signaling. (**A-B**) Elevated NaR cell proliferation in *stc1a^-/-^* fish. Progenies of *stc1a (+17)^+/-^* intercrosses were raised in the E3 embryo medium to 5 dpf. NaR cells were detected by in situ hybridization using an *igfbp5a* cRNA probe. After NaR cells were visualized and quantified in each fish, fish was genotyped individually. Representative images are shown in (**A**) and quantified data in (**B**). Scale bar = 0.2 mm. n = 33-70 larvae/group. **(C**) Increased IGF1 receptor-Akt signaling in *stc1a^-/-^* fish. Progenies of *stc1a (+17)^+/-^; Tg (igfbp5a: GFP)* intercrosses were raised in E3 embryo medium to 3 dpf and treated with DMSO or 0.3 μM BMS-754807 (BMS). Two days later, fish were fixed and phospho-Akt positive cells were detected by immunostaining. These fish were genotyped individually afterwards. n = 14-47 larvae/group. (**DE)** Progeny of *stc1a (+17)^+/-^; Tg (igfbp5a: GFP)* intercrosses were raised in E3 embryo medium and transfer to normal [Ca^2+^] embryo medium containing DMSO, 0.3 μM BMS-754807 (BMS), 0.06 μM Wortmannin (Wort), 8 μM MK2206 (MK), 5 μM Rapamycin (Rapa), or 10 μM U0126 at 3 dpf. Two days later, NaR cells were quantified. They were genotyped individually afterwards. Representative images (**D**) and quantified data are shown (**E**). n = 7-35 larvae/group. Scale bar = 0.2 mm.

We next examined IGF signaling by immunostaining using a Phospho-Akt antibody as reported (Dai et al., 2014). Compared to the siblings, *stc1a (+17)^-/-^* larvae had significantly more phospho-Akt positive cells (Figure 5C). This increase was abolished by the addition of the IGF1 receptor inhibitor BMS-754807 (Figure 5C), suggesting loss of Stc1a increases IGF signaling. Treatment of *stc1a (+17)^-/-^* and *stc1a (Δ18+1)^-/-^* mutant fish with BMS-754807 reduced NaR cell number to the sibling level (Figure 5D-E, Figure 5-figure supplement 1D). Likewise, the PI3 kinase inhibitor wortamannin, Akt inhibitor MK2206, and Tor inhibitor rapamycin all abolished NaR cell proliferation in *stc1a* mutant fish. The addition of the MEK inhibitor U0126 had no such effect level (Figure 5D-E, Figure 5-figure supplement 1D). These data suggest that genetic deletion of *stc1a* increases NaR cell proliferation via activating IGF-PI3 kinase-Akt-Tor signaling.

### Stc1a regulates NaR cell reactivation in a Papp-aa and Igfbp5a-dependent manner

Gene expression analysis showed that *papp-aa* mRNA levels were significantly higher in *stc1a (+17)^-/-^* fish compared with their siblings (Figure 6A). Because mammalian STC1 can bind to and inhibit PAPP-A-mediated IGFBP proteolytic cleavage and since Papp-aa is highly expressed in zebrafish NaR cells (Kløverpris et al., 2015; Liu et al., 2020), we tested whether Stc1a regulates IGF signaling in NaR cells via Papp-aa. Knockdown of Stc1a using validated gRNAs (Figure 6- figure supplemental 1) resulted in a significant increase in NaR cell proliferation in *Tg (igfbp5a: GFP)* fish. This increase was abolished by treatment with ZnCl_2_ or batimastat, two distinct Papp-aa inhibitors (Tallant et al., 2006) (Figure 6B). This increase was also abolished by the IGF1 receptor inhibitor BMS-754807 (Figure 6B), suggesting the involvement of IGF signaling. If Stc1a indeed acts via suppressing Papp-aa-mediated Igfbp5a protelolysis, then loss of Stc1a should not be able to reactivate NaR cells in the absence of Papp-aa or Igfbp5a. Indeed, while knockdown of Stc1a resulted in significant increases in NaR cell proliferation in wild-type and hetergygous siblings, it did not affect NaR cell proliferation in *pappaa^-/-^; Tg (igfbp5a: GFP)* embryos (Figure 6C). Likewise, knockdown of Stc1a in *igfbp5a^-/-^* mutant fish did not increase NaR cell proliferation (Figure 6- figure supplemental 2). This hypothesis were tested further by generating *stc1a^-/-^* and *papp-aa^-/-^* double mutant embryos. As shown in Figure 6D, while permanent deletion of *stc1a* significantly increased NaR cell proliferation, it had no such effect in the *papp-aa^-/-^* mutant background. These data suggest that Stc1a promotes NaR cell quiescence and inhibits NaR cell reactivation by suppressing IGF signaling and this fuction is Papp-aa- and Igfbp5a-dependent.

**Figure 6.**
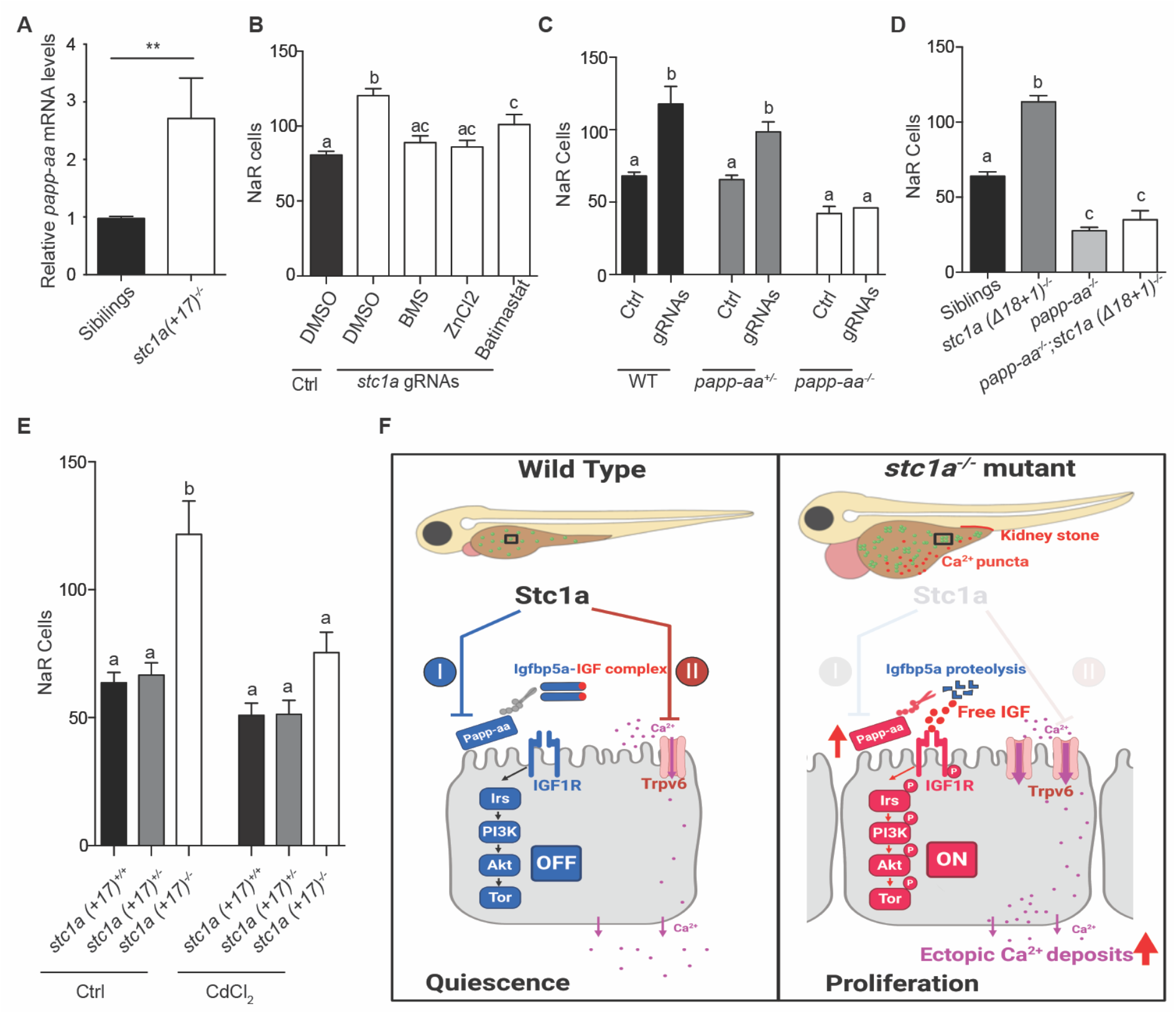
Papp-aa is indispensible in Stc1a regulation of NaR cell quiescence-proliferation balance. (**A**) Embryos of the indicated genotypes were raised in E3 embryo medium until 5 dpf. The mRNA levels of *papp-aa* were measured and normalized. n = 15~17. **, P < 0.001 by unpaired two-tailed t test. **(B)** *Tg(igfbp5a: GFP)* embryos were injected with *stc1a* targeting gRNAs and Cas9 mRNA at the 1-cell stage. Embryos were raised in E3 embryo medium. The injected embryos were treated with BMS-754807 (0.3 μM), ZnCl_2_ (8 μM), or Bastimastat (200 μM) from 3 to 5 dpf. NaR cells were quantified and shown. n= 20~39. (**C**) Progeny of *papp-aa^+/-^; Tg (igfbp5a: GFP)* intercrosses were injected with *stc1a* targeting gRNAs and Cas9 mRNA. The injected embryos were raised and NaR cells were quantified at 5 dpf. Each larva was genotyped afterwards. n= 5~19. **(D)** Larvae of the indicated genotypes were raised in E3 embryo medium and NaR cells were quantified at 5 dpf. n = 9 ~ 78. **(E)** Progeny of *stc1a* (+17)^+/-^*; Tg (igfbp5a: GFP)* intercrosses were raised in E3 embryo medium and treated with or without 10 μg/L CdCl_2_ from 3 dpf to 5 dpf. NaR cells were quantified and shown. Each larva was genotyped afterwards. n = 8-24. **(F)** Proposed model of Stc1a function as a [Ca^2+^]-regulated molecular switch regulating the calcium uptake and epithelial cell quiescence-proliferation balance.

Previous studies have shown that Trpv6 conducts Ca^2+^ influx into NaR cells and promotes NaR cell quiescence by suppresses Akt-Tor signaling (Xin et al., 2019). To determine whether Trpv6 is involved in this action of Stc1a, progenies of *stc1a (+17)^+/-^;Tg (igfbp5a: GFP)* intercrosses were subjected to CdCl_2_ treatment. Inhibition of Trpv6 significantly reduced NaR cell reactivation and proliferation in *stc1a (+17)^-/-^* mutant fish (Figure 6E), suggesting that Trpv6-mediated Ca^2+^ influx is involved in Stc1a’s action in regulating NaR cell quiescence-proliferation balance.

## Discussion

Although Stc1 has been known as a fish hypocalcemic hormone for several decades (Pang, 1973; Gagliardi et al., 2005; Wagner and Dimattia, 2006; Yeung et al., 2012), its longterm in vivo function in fish was not clear due to the lack of a genetic mutant. Despite many years of investigations, how Stc1 acts in vivo is not well understood and the molecular identity of Stc1 receptor(s) is still unknown. Moreover, most if not all vertebrates have multiple *stc* genes. Whether these genes have similar functions or act redundantly is not clear. In this study, we generated several *stc* mutants and show Stc1a not only inhibits Ca^2+^ uptake via regulating Trpv6 expression and activity, but also play a previously unrecognized role regulating epithelial cell quiescence-proliferation decision. Furthermore, we have delineated a [Ca^2+^]-regulated pathway turns on/off local IGF activity in epithelial cells locally.

The *stc1a^-/-^* mutant fish had normal growth, developmental speed, and morphology during early development. At 4 dpf, however, *stc1a^-/-^* mutant fish began to display cardiac edema and this was followed by body swelling from 5-7 dpf. Additionally, *stc1a-/-* fish had no inflated swim bladder and showed impaired touch and swim response. All mutant fish died by 10 dpf. These phenotypes are specific since the *stc2b^-/-^* mutant fish did not exhibit such phenotypes. Biochemical analysis results show that loss of Stc1a increased total body calcium levels. Moreover, *stc1a^-/-^* mutant fish developed ectopic calcification on the yolk sac region and in the kidney. Mechanistic analysis suggested that Stc1a regulates the body Ca^2+^ balance by reducing Trpv6 expression and Trpv6-mediated Ca^2+^ uptake. This conclusion is supported by several lines of evidence. First, the expression of *trpv6* mRNA is significantly elevated in *stc1a^-/-^* mutant fish. Second, pharmacological inhibition of Trpv6 markedly in *stc1a^-/-^* mutant fish reduced ectopic calcification and kidney stone formation. Trpv6 inhibition also abolished cardiac edema and body swelling and rescued *stc1a^-/-^* mutant fish from premature death. Third, Ca^2+^ depletion reduced ectopic calfication and kidney stone formation and prevented the premature death in the mutant fish. Our loss-of-function genetic study results agree well with classical physiology studies. Surgical removal of CS resulted in the development of kidney stones in killifish (Pang, 1971). Injection CS extracts or purified Stc1 reduced epithelial Ca^2+^ uptake in killifish, salmon, eel, carp, and other fish (Pang, 1973). In zebrafish, overexpression of Stc1a by mRNA injection into the embryos reduced Ca^2+^ uptake and decreased *trpv6* mRNA expression (Tseng et al., 2009). The *stc1a^-/-^* mutant fish phenotypes reported in this study, however, differ considerably from those reported in the mouse model. Stc1-null mice grew and reproduced normally and no anatomical or histological abnormalities were detected in adult tissues (Chang et al., 2005). There are several possible explanations for the differences. Mammals do not have CS glands or comparable structure. In mammals, *STC1* gene is expressed in a wide range of tissues and acts mainly as a paracrine and autocrine factor (Yeung et al., 2012). Published evidence suggests that STC-1 does not appear to play a major role in calcium homeostasis regulation (Wagner and Dimattia, 2006; Yeung et al., 2012). Unlike mammals, freshwater teleost such as zebrafish live in hypoosmotic aquatic habitats. Their body fluids are hyperosmotic compared to the surrounding water (Evans, 2008). To maintain body osmolarity, they constantly remove osmotic water by secreting a large volume of diluted urine from the kidney (Evans, 2008). Therefore, the development of cardiac edema and subsequently body swelling in *stc1a* mutant fish may be resulted from the accumulation of osmotic water, caused by defective kidney functions and/or an increased Ca^2+^ uptake and elevated body Ca^2+^ levels. This speculation is supported by the fact that inhibition of Trpv6-mediated Ca^2+^ uptake not only decreased ectopic Ca^2+^ deposit and kidney stones, but also prevented the development of cardiac edema and body swelling and premature death. Likewise, depletion of Ca^2+^ alleviated the edema and body swelling and rescued fish from premature death.

A new and important finding made in this study is that Stc1a functions as a [Ca^2+^]- dependent regulator of epithelial cell quiescence. The normally quiescent NaR cells re-enter the cell cycle in response to low Ca^2+^ stress and the NaR cell reactivation has been attributed to the activation of local IGF signaling (Dai et al., 2014; Liu et al., 2017; Liu et al., 2018). NaR cell reactivation is impaired in *igfbp5a^-/-^* mutant fish (Liu et al., 2018). Likewise, genetic deletion and pharmacological inhibition of Papp-aa, the Igfbp5a proteinase, abolishes NaR cell reactivation and IGF signaling (Liu et al., 2020). The Papp-aa-mediated Igfbp5a proteolysis is suppressed by a [Ca^2+^]-dependent mechanism (Liu et al., 2020). Although previous studies have shown that Stc1 expression is increased in fish raised in high [Ca^2+^] media (Kwong et al., 2014; Lin et al., 2014; Radman et al., 2002; Tseng et al., 2009), less is known about the other *stc* genes. Whether the expression of *stc* genes is increased by body Ca^2+^ states is unclear. In this study, we investigated the impact of Ca^2+^ states on the 4 *stc* genes in zebrafish. In addition of manipulating [Ca^2+^] in the media as an indirect way to alter body Ca^2+^ levels, we also utilized *trpv6^-/-^* and *papp-aa^-/-^* mutant fish, which have been shown to be hypocalcemic (Liu et al., 2020; Xin et al., 2019). Our data suggest that the expression of *stc1a*, but not other *stc* genes, is regulated in a [Ca^2+^]-dependent manner. Genetic deletion of *stc1a*, but not *stc2b*, led to NaR cell reactivation. These findings suggest that high levels of Stc1a promotes NaR cell quiescence under [Ca^2+^] rich conditions. This action is clearly mediated via IGF signaling. Loss of Stc1a resulted in the activation of IGF1 receptor-mediated Akt signaling in NaR cells. Pharmacological inhibition of IGF1 receptor, PI3 kinase, Akt, and Tor all abolished NaR cell reactivation in *stc1a^-/-^* mutant fish. In contrast, Mek-Erk inhibition did not have such effect, suggesting the MAP kinase pathway does not play a major role. In further support of our conclusion, loss of Stc1a did not reactivate NaR cells in *papp-aa^-/-^* mutants, while it did so in the wild-type and heterozygous siblings. Likewise, loss of Stc1a did not increase NaR cell proliferation *igfbp5a^-/-^* fish. These findings, together with our previous studies (Liu et al., 2020; Liu et al., 2018), suggest that under [Ca^2+^] rich conditions, Stc1a is highly expressed and it inhibits Papp-aa-mediated Igfbp5a proteolysis. The intact Igfbp5a binds to and sequesters IGFs in the Igfbp5a/IGF complex. This turns off IGF signaling in NaR cells and keeps these cells in the quiescent state (Fig. 6F). Under low [Ca^2+^] conditions or in *stc1a^-/-^* fish, the Stc1a levels are reduced and Papp-a-mediated Igfbp5a proteolysis increases. This liberates IGFs from the Igfbp5a/IGF complex. The free IGFs bind to the IGF1 receptor and turns on IGF signaling and promotes NaR cells to re-enter the cell cycle and proliferate (Fig. 6F).

Although our conclusion is based on findings made in zebrafish, available evidence suggests that this regulatory loop is likely operational in mammals. Human STC1 can bind to PAPP-A and inhibit PAPP-A proteinase activity (Kløverpris et al., 2015). This activity is conserved from fish to humans (Liu et al., 2020). Human PAPP-A primarily cleaves IGFBP4 and IGFBP5 (Conover and Oxvig, 2018; Laursen et al., 2007; Oxvig, 2015). A number of studies indicate a local role of STC1 in the kidney (Kashyap et al., 2020b). Genome-wide association studies have identified STC1 as one of the chronic kidney disease genes (Böger and Heid, 2011). Treatment with human STC1 decreased gluconeogenesis in rat renal medulla slices and in fish kidney slices (Schein et al., 2012). While global deletion of STC1 in C57BL/6 mice did not cause major abnormalities (Chang et al., 2005), a recent study using humanized ex vivo and in vivo mouse models suggested that mesenchymal stromal cells secrete STC1 to suppress hematopoietic stem cell (HSC) proliferation (Waclawiczek et al., 2020). Another recent study reported that renal expression of STC1, IGF-1, IGF1 receptor, IGFBP5, and PAPP-A are all increased in an autosomal dominant polycystic disease (ADPKD) mouse model (Kashyap et al., 2020a). STC1 knockout significantly decreased cyst development and improved kidney injury in the ADPKD mice (Kashyap et al., 2020a).

In summary, this study has shown that Stc1a plays two roles in regulating calcium homeostasis and cell quiescence balance and implicates Trpv6-mediated Ca^2+^ uptake and Papp-aa-Igfbp5a-mediated IGF signaling in these two functions (Figure 6F). The Trpv6-mediated Ca^2+^ uptake and IGF signaling are likely linked. In a recent study, we have shown that Trpv6-mediated constitutive Ca^2+^ influx promotes cell quiescence state by suppressing IGF signaling (Xin et al., 2019). In this study, we found that the elevated NaR cell reactivation in the *stc1a^-/-^* mutant fish requires Trpv6. These findings not only provide genetic evidence and mechanistic insights into the hypocalcemic function of Stc1a, but also provided new insights into cell quiescence regulation. Although insulin/IGF-PI3 kinase-Akt-Tor signaling has been implicated in adult stem cell reactivation in fly, mouse, and zebrafish, the underlying mechanisms are still not well understood. The discovery made in this study, together with a recent report that STC1 inhibits HSC proliferation in mice (Waclawiczek et al., 2020) has raised the question of whether STC1/Stc1 plays a critical role in reactivating adult stem cells by turning off IGF signaling in these cells. The genetic analysis results in this study show that Stc1a regulates NaR cell reactivation in a Papp-aa-dependent manner. Papp-aa is highly expressed in NaR cells (Liu et al., 2020). Mammalian PAPP-A has five short consensus repeats (SCRs) that are involved in the binding to proteoglycans on the cell surface (Laursen et al., 2002). These structural motifs are conserved in zebrafish (Wolman et al., 2015). It is possible that zebrafish Papp-aa is tethered to the cell-surface and binds to Stc1a, as the case with human PAPP-A (Jepsen et al., 2015; Kløverpris et al., 2015). Future studies are needed to test whether Papp-aa functions as a Stc1a receptor in fish and in mammals.

## Materials and methods

### key resources table

#### Chemicals and reagents

Chemical and molecular biology reagents were purchased from Fisher Scientific (Pittsburgh, PA) unless otherwise noted. Alizarin Red S, ZnCl_2_, batimastat and GdCl_3_ were purchased from Sigma (St. Louis, MO, USA), MK2206 from ChemieTek (Indianapolis, IN) and Rapamycin from Calbiochem (Gibbstown, NJ). BMS-754807 was purchased from Active Biochemicals Co.. The Phospho-Akt antibody was purchased from Cell Signaling Technology (Danvers, MA, USA) and various restriction enzymes were purchased from New England BioLabs (Ipswich, MA, USA). Primers, TRIzol, M-MLV reverse transcriptase were purchased from Life Technologies (Carlsbad, CA, USA). Anti-Digoxigenin-AP antibodies was purchased from Roche (Basel, Switzerland). The pT3.Cas 9-UTRglobin vector was a gift from Dr. Yonghua Sun, Institute of Hydrobiology, Chinese Academy Sciences.

#### Zebrafish

Zebrafish were maintained, crossed, and staged in accordance to standard guidelines (Kimmel et al., 1995; Westerfield, 2000). Embryos and larvae were raised at 28.5°C in standard E3 embryo medium. Three additional embryo rearing solutions, referred as normal [Ca^2+^] medium (containing 0.2 mM [Ca^2+^]), low [Ca^2+^] medium (containing 0.02 mM[Ca^2+^]) and very low [Ca^2+^] medium (containing 0.001 mM [Ca^2+^]), were prepared as previously reported (Dai et al., 2014). To inhibit pigmentation, 0.003% (w/v) N-phenylthiourea was added to these solutions beginning at 20–22 hours post fertilization (hpf). *Tg(igfbp5a:GFP), trpv6^-+/-^*, and *igfbp5a^-/-^* fish lines were generated in previous studies (Liu et al., 2020; Liu et al., 2018; Xin et al., 2019). The *stc2b^+/-^* fish (*stc2b^sa24026^*, ZIRC# ZL10776) were obtained from ZIRC. All experiments were conducted in accordance with the guidelines approved by the Institutional Committee on the Use and Care of Animals, University of Michigan.

#### RT-qPCR

Total RNA was extracted from pooled zebrafish embryos and larvae. RNA was reverse-transcribed to cDNA using oligo(dT)18 primer and M-MLV (Promega). qPCR was performed using SYBR Green (Bio-Rad) on a StepONE PLUS real-time thermocycler (Applied Biosystems). PCR primers were designed based on the 4 zebrafish *stc* gene sequences obtained from the Ensemble database (Ensemble gene numbers are: *stc1a*, ENSDARG00000058476, *stc1b*, ENSDARG00000003303, *stc2a*, ENSDARG00000056680, and *stc2b*, ENSDARG00000102206). The expression level of a target gene transcript was normalized by *β-actin* or *18S* RNA level. The following primers used are: *stc1a*-qPCR-F: 5’-CCAGCTGCTTCAAAACAAACC-3’, *stc1a*-qPCR-R: 5’-ATGGAGCGTTTTCTGGCGA-3’, *stc1b*-qPCR-F: 5’-CCAAGCCACTTTCCCAACAG-3’, *stc1b*-qPCR-R: 5’-ACCCACCACGAGTCTCCATTC-3’, *stc2a*-qPCR-F: 5’-TATGGTCTTCCAGCTTCAGCG-3’, *stc2a*-qPCR-R: 5’-CGAGTAATGGCTTCCTTCACCT-3’, *stc2b*-qPCR-F: 5’-CACAAGAAAAGACTGTCTCTGCAGA-3’, *stc2b*-qPCR-R: 5’-GGTAGTGACATCTGGGACGG-3’, *trpv6*-qPCR-F: 5’ - GGACCCTACGTCATTGTGATAC-3’, *trpv6*-qPCR-R: 5’-GGTACTGCGGAAGTGCTAAG-3’, *papp-aa*-qPCR-F: 5’-AAAGAGGAGGGCGTTCAAG-3’, *papp-aa*-qPCR-R: 5’-TGCAGCGGATCACATTAGAG-3’(Liu et al., 2020), *18s*-qPCR-F: 5’-AATCGCATTTGCCATCACCG-3’, *18s*-qPCR-R: 5’-TCACCACCCTCTCAACCTCA-3’, *β-actin*-qPCR-F: 5’-GATCTGGCATCACACCTTCTAC-3’, *β-actin*-qPCR-R: 5’-CCTGGATGGCCACATACAT-3’.

#### Generation of *stc1a^-/-^* fish lines and transient knockdown of *stc1a* by CRISPR/Cas9

Two sgRNAs targeting the *stc1a* gene were designed using CHOPCHOP (http://chopchop.cbu.uib.no). Their sequences are: *stc1a*-sgRNA-1, 5’-GCAGAGCGCCATTCAGACAG-3’ and *stc1a*-sgRNA-2, 5’-GCAGATCTCGTGCATGCCGT-3’. Mixed sgRNA (30-40 ng/μl) and Cas 9 mRNA (200-400 ng/μl) were injected into *Tg(igfbp5a:GFP)* embryos at the 1-cell stage (Xin and Duan, 2018). A subset of injected F0 embryos was used to identify indels. DNA was isolated from individual embryo and analyzed by PCR followed by hetero-duplex assays as reported (Liu et al., 2018). For transient knockout experiments, the remaining injected F0 embryos was raised in E3 embryo medium to 3 dpf and transferred to the intended embryo medium from 3 to 5 dpf as previously reported (Dai et al., 2014). To generate stable *stc1a^-/-^* fish lines, injected F0 embryos were raised to adulthood and crossed with *Tg(igfbp5a:GFP)* or wild-type fish. F1 fish was raised to adulthood and genotyped. After confirming indels by DNA sequencing, the heterozygous F1 fish were intercrossed to generate F2 fish.

#### Genotyping

Genomic DNA was isolated from individual or pooled fish as reported (Liu et al., 2018). HRMA was performed to genotype *igfbp5a-/-, trpv6-/-* fish, *papp-aa-/-* fish and siblings following published methods (Liu et al., 2020; Liu et al., 2018; Xin et al., 2019). The *stc1a^+/-^* mutant fish genotyping was performed by PCR using the following primers: *stc1a*-gt-F, 5’-TGAAAACCACTGCCTTAAATTG-3’*, stc1a*-gt-R, 5’-GTAGCTCTACCGATCCCAAATG-3’. The progenies of *stc2b^+/-^* intercrosses were genotyped by direct DNA sequencing.

#### Morphology analysis

Body length, somite number and head trunk angles were measured as described (Kamei et al., 2011; Liu et al., 2018). Alizarin red staining was conducted as previously reported (Du et al., 2001). The bright-field images were acquired using a stereomicroscope (Leica MZ16F, Leica, Wetzlar, Germany) equipped with a QImaging QICAM camera (QImaging, Surrey, BC, Canada).

#### Total body Ca^2+^ assay

The sample preparation was carried out as described by (Elizondo et al., 2010). Briefly, fish larvae were anesthetized using MS-222. For each sample, 30 embryos were pooled and washed twice with deionized water, and dried at 65 °C for 60 min. Next, 125 μL 1M HCl was added to each tube and incubated overnight at 95 °C with shaking. Next, samples were centrifuged and the supernatant was collected. Total calcium content was measured using a commercial kit (ab102505, Abcam, USA) following the manufacturer’s instruction.

#### Live imaging and microscopy

NaR cells were quantified as previously reported (Liu et al., 2017). Briefly, embryos and larvae were anesthetized with MS-222. Larvae were mounted and subjected to fluorescent imaging using the Leica MZ16F stereo microscope. Image J was used for image analysis and data quantification.

#### Immunostaining and whole mount in situ hybridization

Whole mount immunostaining and in situ hybridization were performed as described previously (Dai et al., 2014). Briefly, samples were fixed in 4% paraformaldehyde and permeabilized in methanol. Larvae were incubated overnight with the phospho-Akt antibody in 4°C, followed by incubation with an anti-rabbit HRP antibody (Jackson ImmunoResearch, West Grove, PA, USA) and by nickel-diaminobenzidine staining. For in situ hybridization, *igfbp5a* mRNA signal was detected using a digoxigenin (DIG)-labeled antisense riboprobe. Larvae were incubated in an anti-DIG-AP antibody (Roche) and development in BCIP-NBT staining buffer prior to imaging and analysis.

#### Drug treatment

All drugs except ZnCl_2_ and CdCl_2_ used in this study, were dissolved in DMSO and further diluted in double-distilled water. ZnCl_2_ was dissolved in distilled water. Zebrafish larvae were treated with ZnCl_2_, batimastat, BMS-754807, and other drugs from 3dpf to 5 pdf (Dai et al., 2014; Liu et al., 2017). Drug solutions were changed daily. 5dpf larvae were then collected for immunostaining, in situ hybridization or fluorescent imaging.

#### Statistical analysis

Statistical tests were carried out using GraphPad Prism 8 software (GraphPad Software, Inc., San Diego, CA). Values are shown as means ± SEM. Statistical significance between experimental groups was performed using unpaired two-tailed t test, Chi-square test, long-rank test and one-way ANOVA followed by Tukey’s multiple comparison test. Statistical significances were accepted at *P* < 0.05 or greater.

## Acknowledgments

This work was supported by NSF grant IOS-1557850 and University of Michigan M-Cubed 3 Project U064122 to CD. The funders had no role in study design, data collection and analysis, decision to publish, or preparation of the manuscript.

## Supplementary materials for this manuscript include the following

### Supplemental figure 1-14

**Figure 1- Supplemental Figure 1.**
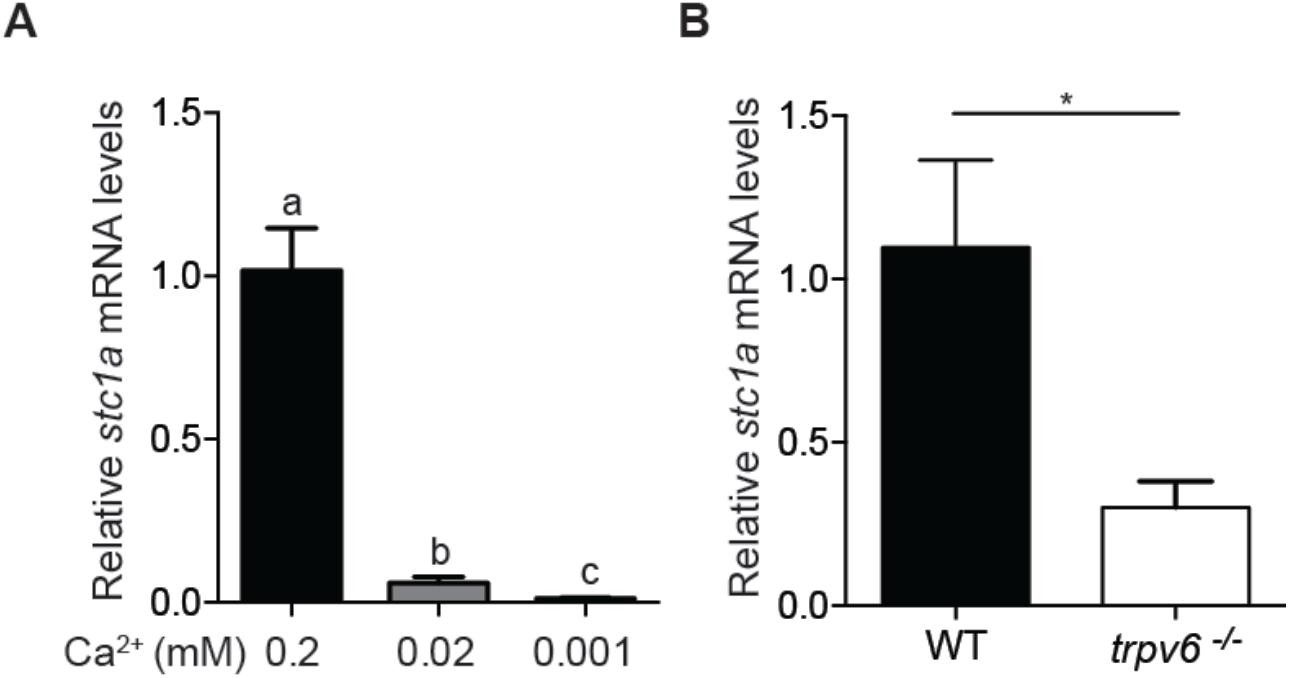
Regulation of *stc1a* expression by Ca^2+^ states in larval zebrafish. (A) Wild-type zebrafish were raised in embryo media containing the indicated Ca^2+^ concentration from 5 to 7 dpf. The mRNA expression levels of *stc1a* were determined by qPCR and normalized by β-actin mRNA levels. (B) Zebrafish embryos of the indicated genotypes were raised in E3 embryo medium until 5 dpf. The level of *stc1a mRNA* was measured and shown. Data shown are from 3 independent experiments, each containing 10-15 larvae/group. *, *P* < 0.05 by unpaired two-tailed t test.

**Figure 2- Supplemental Figure 1.**
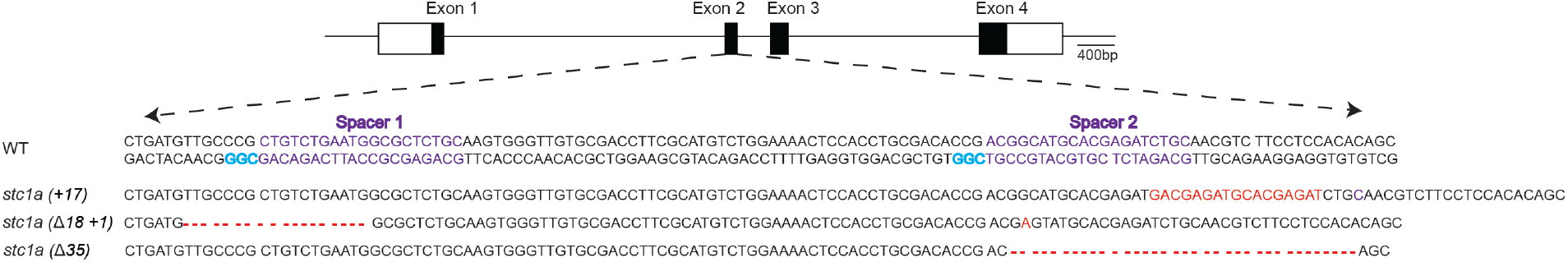
Schematic diagram of *stc1a* and the CRISPR/Cas9 targeting sites. Exons are represented as boxes and introns as lines. Open and filled boxes are untranslated and protein coding regions, respectively. The DNA sequence of wild-type fish (WT) and the indicated mutants are shown. The PAM motif is highlighted in blue. The deletion or insertion is highlighted in red.

**Figure 2- Supplemental Figure 2.**
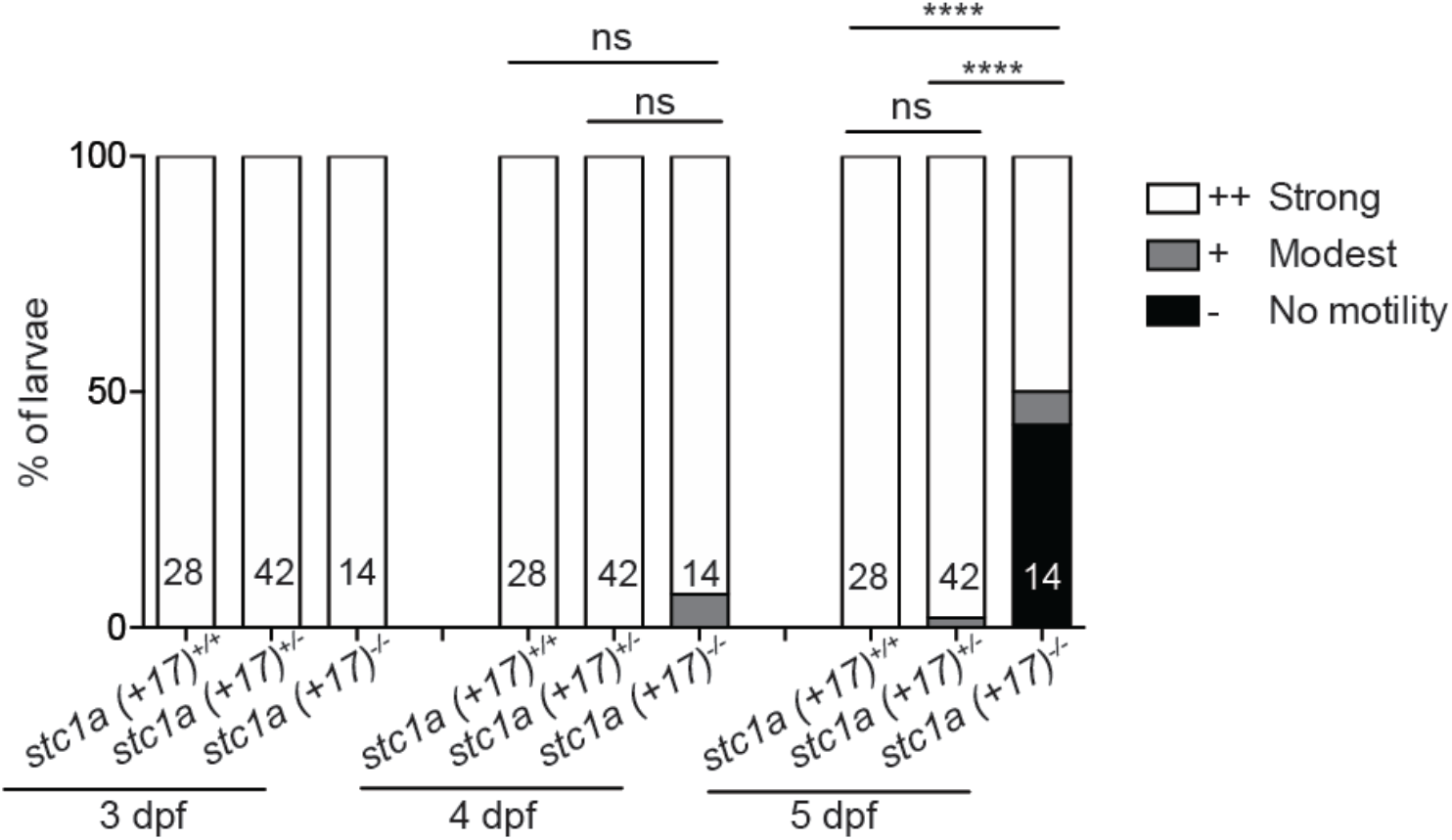
Genetic deletion of *stc1a* impairs the touch-swim response. Progenies of *stc1a* (+17)^+/-^ intercrosses were raised in E3 embryo medium to the indicated stages. Each fish was touched by a pair of tweezers and scored for no motility, modest motility and strong motility. They were genotyped individually afterwards. Number of fish are shown in the column. ****, P< 0.001 by Chi-square test.

**Figure 2- Supplemental Figure 3.**
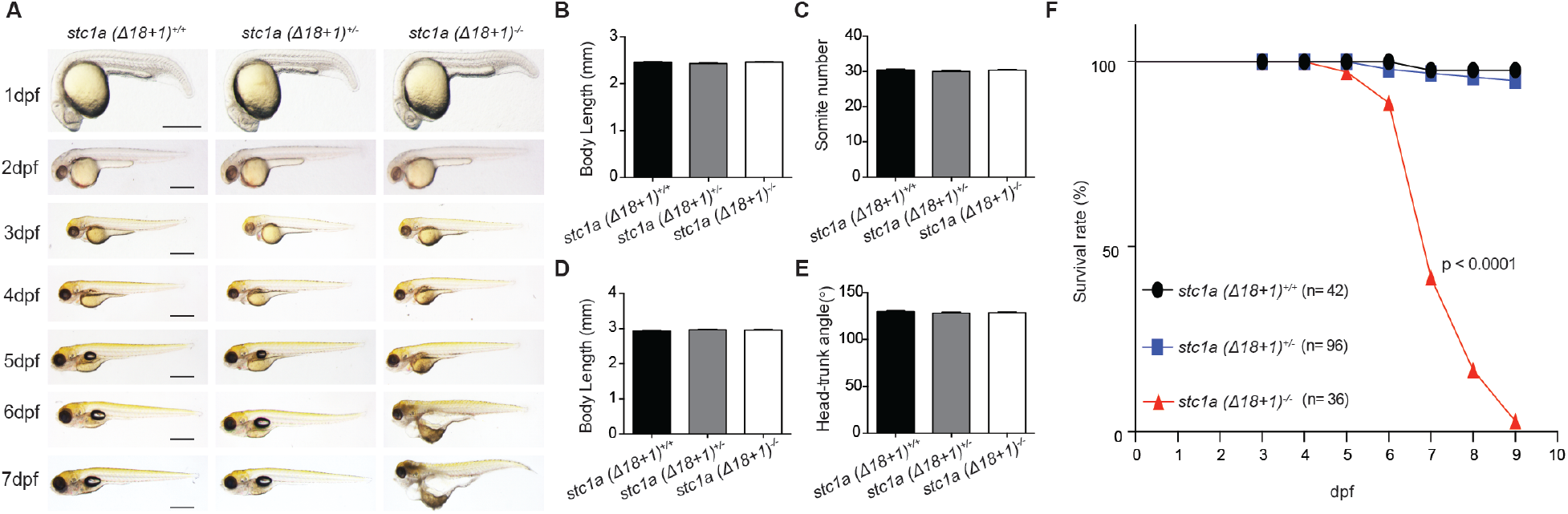
Genetic deletion of Stc1a results in cardiac edema, body swelling, and premature death. (A) Gross morphology of wild-type (WT), *stc1a (Δ18+1)^+/-^* and *stc1a (Δ18+1)^-/-^* fish at the indicated stages. Representative images are shown. Scale bar = 0.5 mm. (B-E) Body length, somite number, and head trunk angle of wild-type (WT), *stc1a (Δ18+1)^+/-^* and *stc1a (Δ18+1)^-/-^* fish were measured at 24 hpf (B, C) and 48 hpf (D, E). n = 14-39 larvae/group. (F) Survival curves. Progeny of *stc1a (Δ18+1)^+/-^* intercrosses were raised in E3 embryo medium. Dead embryos were collected daily and genotyped individually. The survival curves of indicated genotypes and the total fish numbers are shown. *P* < 0.0001 by log-rank test.

**Figure 2- Supplemental Figure 4.**
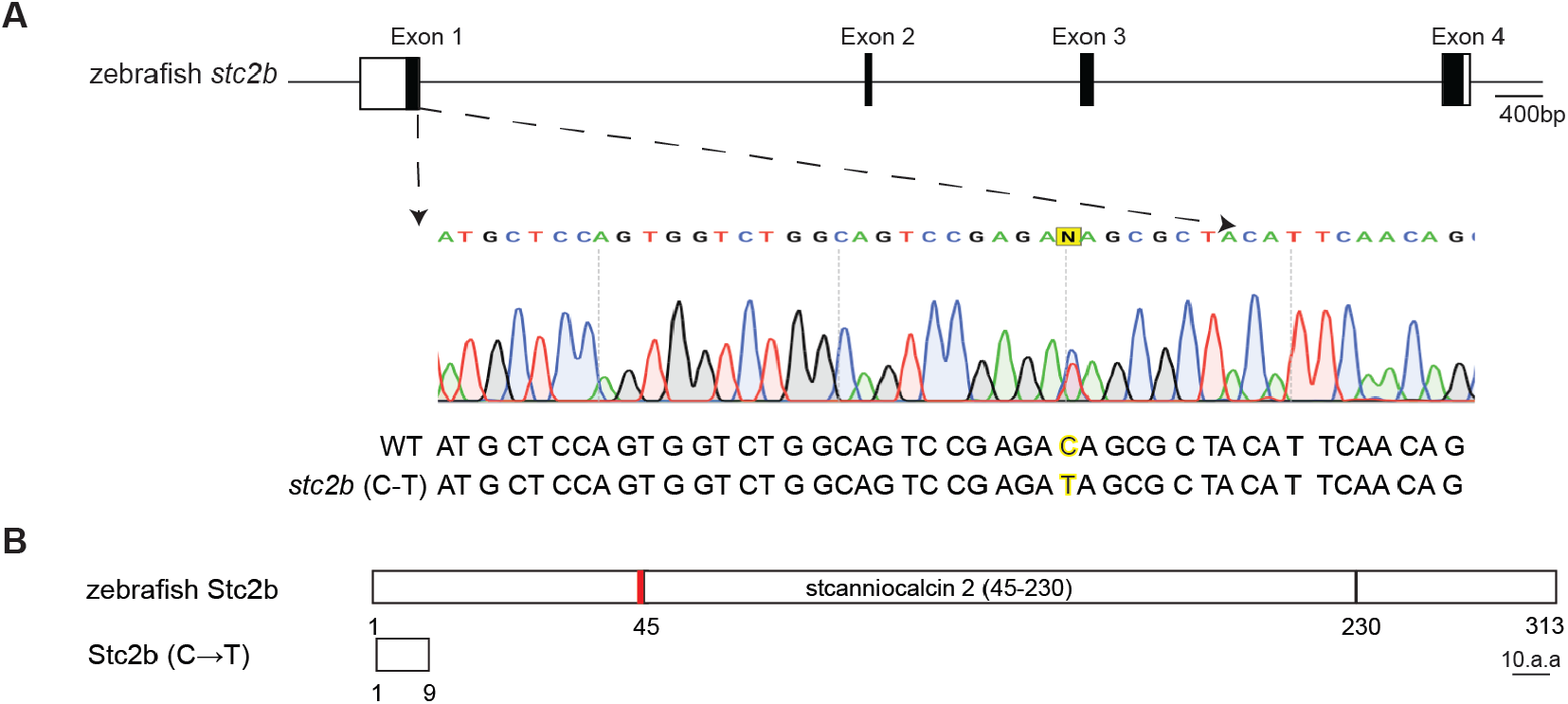
(A) Schematic diagram of the *stc2b* and *stc2b^-/-^* mutant sequence. Exons are represented as boxes and introns as lines. Open and filled boxes are untranslated and protein coding regions, respectively. DNA sequence of *stc2b* and *stc2b^-/-^*. The altered base is shown in yellow. (B) Schematic diagram of Stc2b protein and its mutants. The N-linked glycosylation site is shown by the red bar.

**Figure 2- Supplemental Figure 5.**
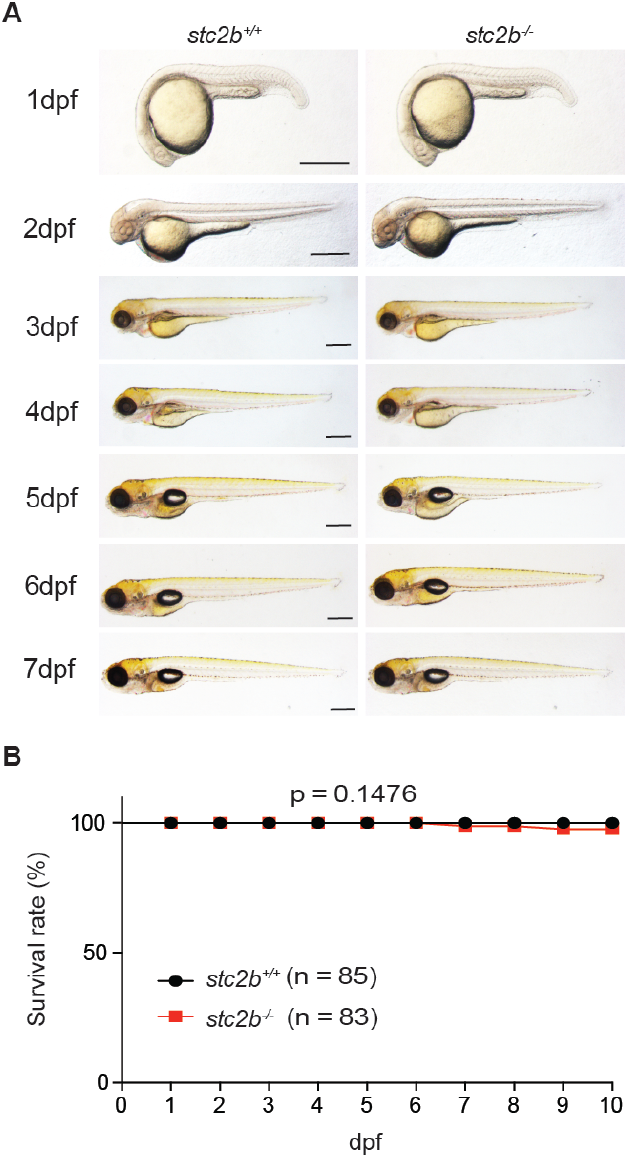
(A) Gross morphology of wild-type and *stc2b^-/-^* fish at the indicated stages. Representative images are shown. Scale bar = 0.5 mm. (B) Survival curves of wild-type (*stc2b^+/+^*) and *stc2b^-/-^* fish. The total fish numbers are shown.

**Figure 3- Supplemental Figure 1.**
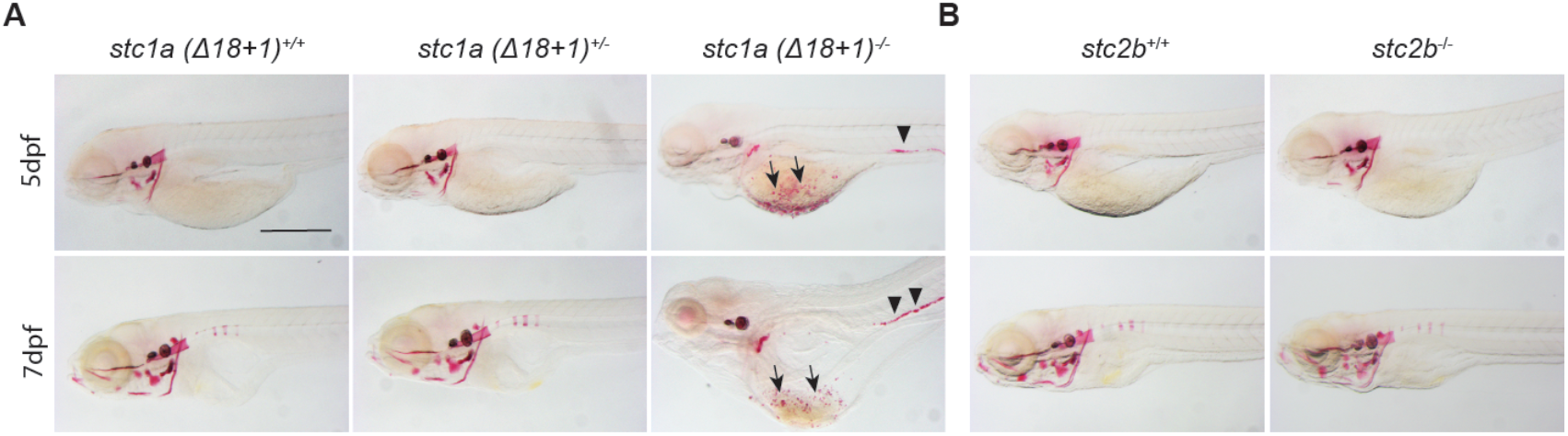
(A-B) Fish of the indicated genotypes were raised in E3 embryo medium and subjected to Alizarin red staining at 5 dpf and 7 dpf. Representative images are shown. Scale bar = 0.5 mm. Note the ectopic calcification in the yolk sac region (arrow) and kidney (arrow heads) in the *stc1a (Δ18+1)^-/-^* mutant fish.

**Figure 3- Supplemental Figure 2.**
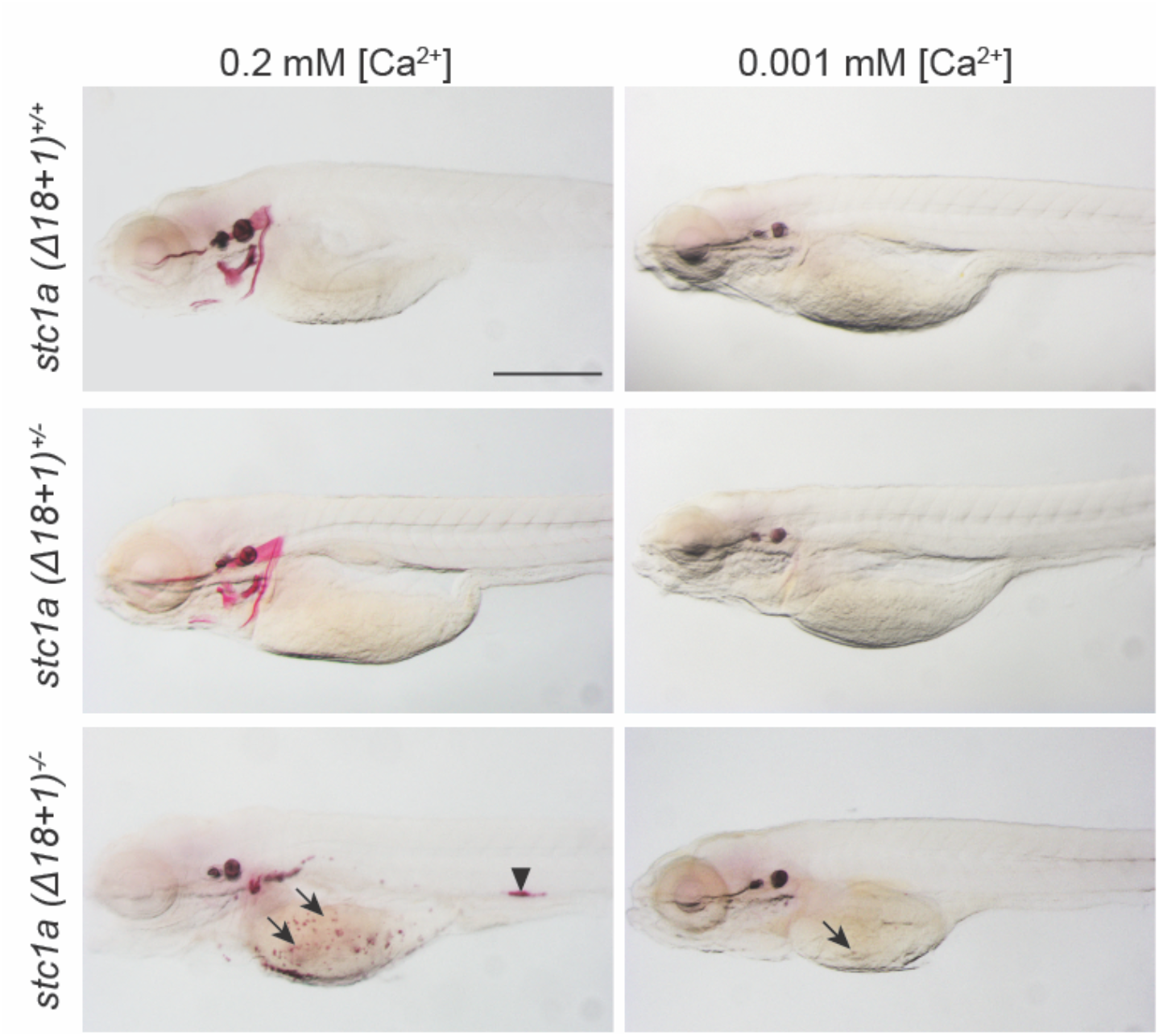
Ca^2+^ depletion reduces ectopic calcification and kidney stone formation. Progenies of *stc1a (Δ18+1)^+/-^* intercrosses were raised in E3 medium to 3 dpf and transferred to the low [Ca^2+^] (0.001 mM) or the control (0.2 mM) embryo medium. Two days later, the larvae were subjected to Alizarin red staining. Each fish was genotyped afterwards. Scale bar = 0.5 mm. Note the ectopic calcification in the yolk sac region (arrow) and kidney stones (arrow heads) in the mutant fish.

**Figure 3- Supplemental Figure 3.**
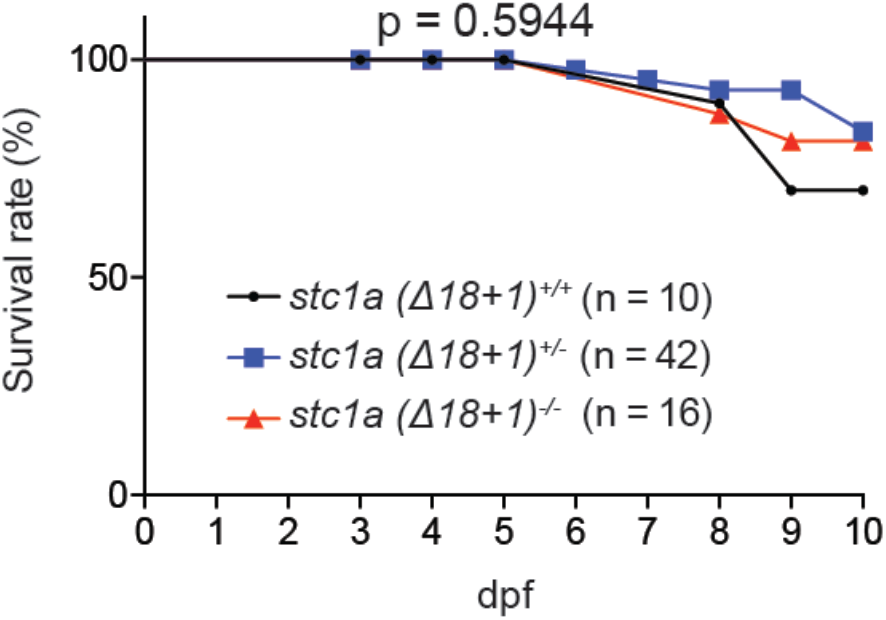
Survival curves of fish of the indicated genotype under low [Ca^2+^]. Progeny of *stc1a (Δ18+1)*^+/-^ intercrosses were raised in the E3 embryo medium till 3 dpf. They were then transferred to the low [Ca^2+^] embryo medium and raised to 10 dpf. Dead fish were collected daily and genotyped individually. No statistical significance was detected among the three genotypes by log-rank test.

**Figure 4- Supplemental Figure 1.**
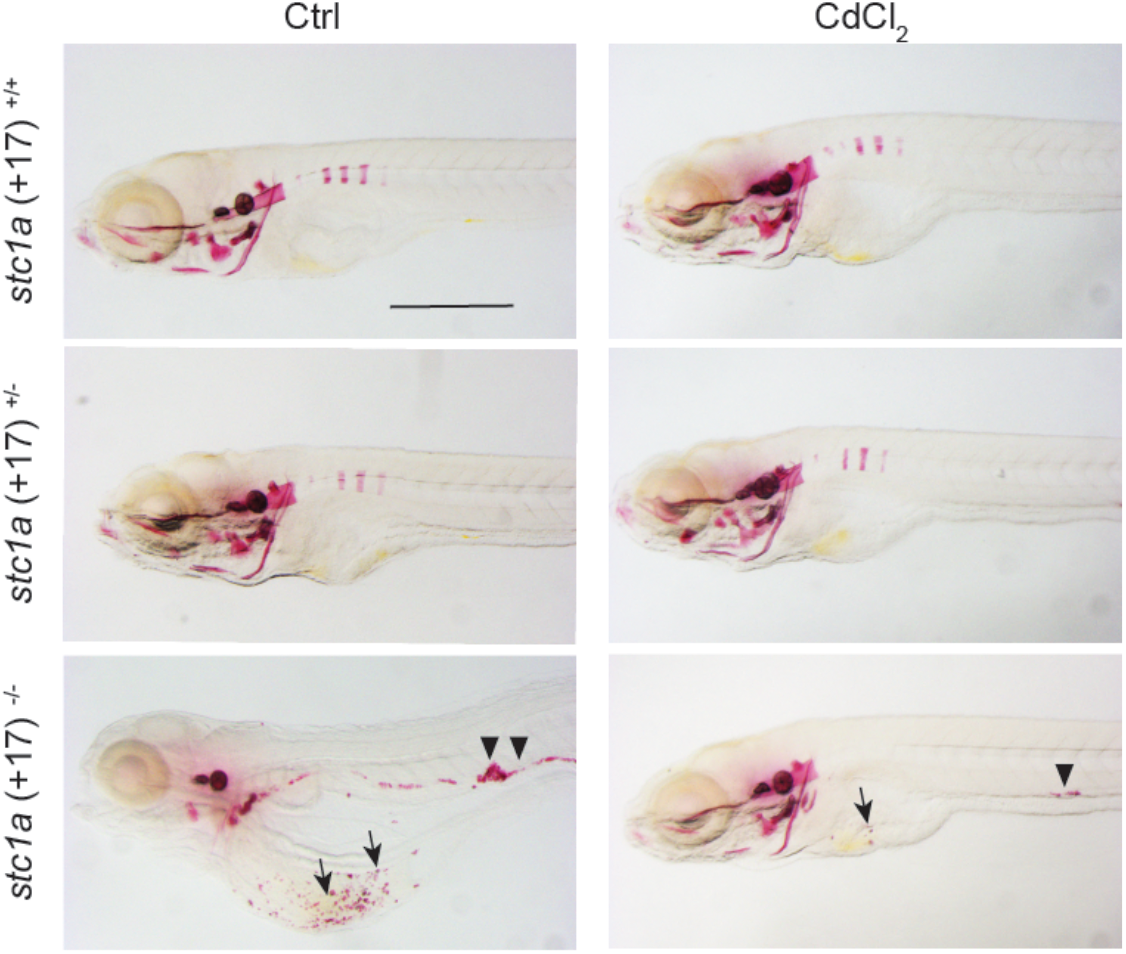
Inhibition of Trpv6 by CdCl_2_ reduces ectopic calcification and kidney stone formation. Progeny of *stc1a (+17)^+/-^* intercrosses were raised in E3 embryo medium and treated with DMSO or 100 μg/L CdCl_2_ from 3 to 7 dpf. The treated larvae were subjected to Alizarin red staining. Each fish was genotyped afterwards. Scale bar = 0.5 mm. Note the ectopic calcification in the yolk sac region (arrow) and kidney stones (arrow heads) in the mutant fish.

**Figure 4- Supplemental Figure 2.**
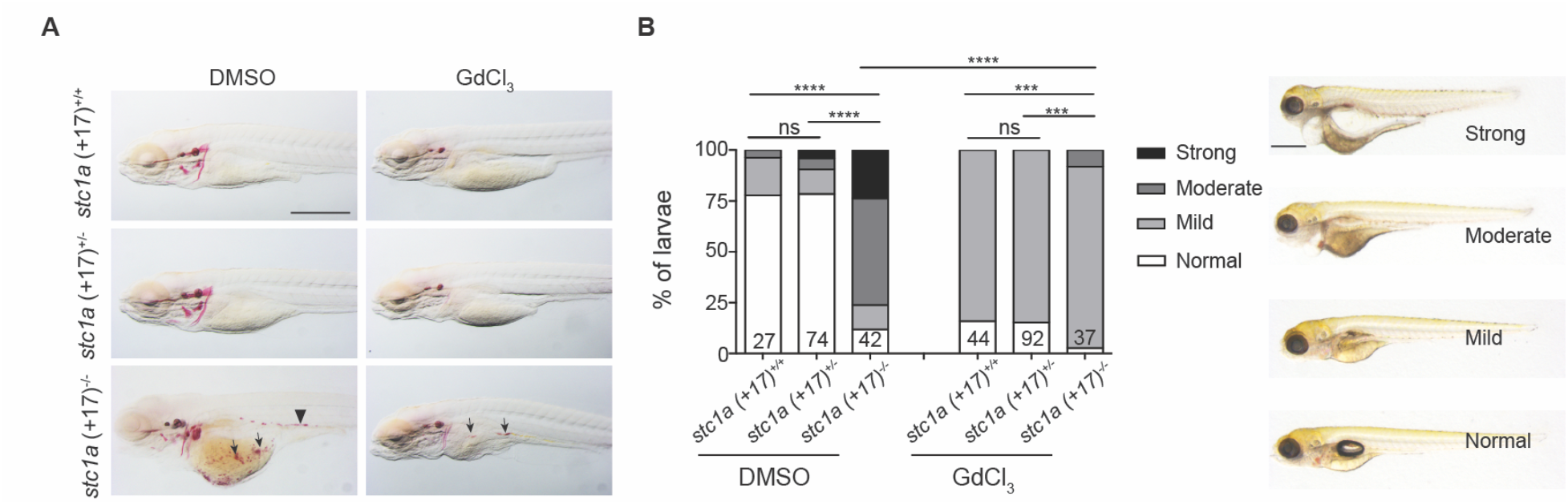
Inhibition of Trpv6 by GdCl_3_ reduces ectopic calcification and kidney stone formation. (A) Progeny of *stc1a (+17)^+/-^* intercrosses were raised in E3 embryo medium and treated with DMSO or 100 μM GdCl_3_ at 3 dpf. At 5 dpf, the treated larvae were subjected to Alizarin red staining. Each fish was genotyped afterwards. Scale bar = 0.5 mm. Note the ectopic calcification in the yolk sac region (arrow) and kidney stones (arrow heads) in the mutant fish. (B) Inhibition of Trpv6 by GdCl_3_ alleviates the edema and swelling phenotypes in *stc1a^-/-^* mutants. Progenies of and *stc1a (+17)^+/-^* intercrosses were treated as described in (A). The fish were scored based on phenotype categories shown on the left. The number of fish in each group is shown in the column. *** and \*\*\*\**P* < 0.01 and <0.001 by *Chi*-square test. Fish numbers are indicated in each column.

**Figure 5- Supplemental Figure 1.**
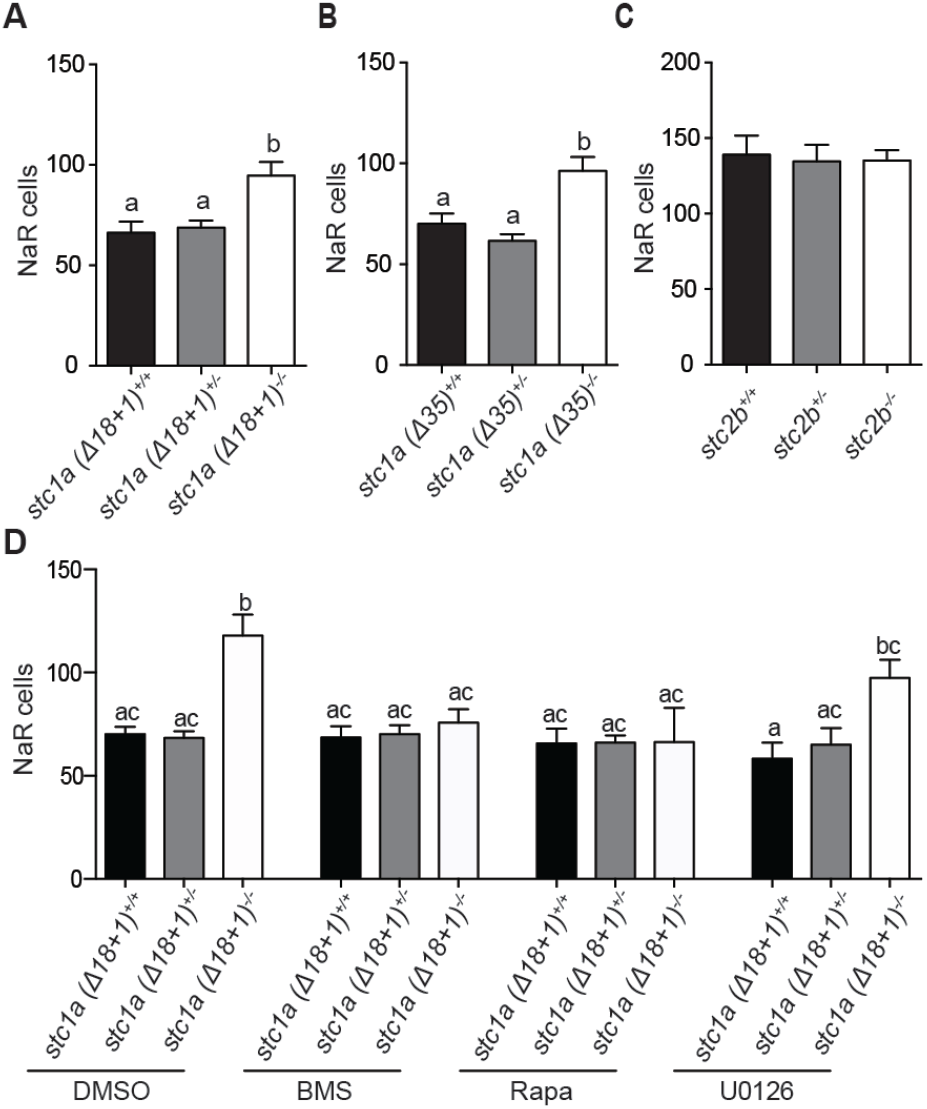
Loss of Stc1a but not Stc2b reactivates NaR cells in an IGF signaling-dependent manner. (A-B) Progeny of *stc1a (Δ18+1)^+/-^; Tg (igfbp5a: GFP)* intercrosses (A), *stc1a (Δ35)^+/-^; Tg (igfbp5a: GFP)* intercrosses (B), or of *stc2b^+/-;^ (igfbp5a: GFP)* intercrosses (C) were raised and NaR cells quantified as described in Figure D. n = 6 - 48 larvae/group. (D) Progeny of *stc1a (Δ18+1)^+/-^; Tg (igfbp5a: GFP)* intercrosses were raised in E3 embryo medium to 3 dpf and transferred to normal [Ca^2+^] embryo medium containing DMSO, 0.3 μM BMS-754807 (BMS), 5 μM Rapamycin (Rapa), or 10 μM U0126. At 5 dpf, NaR cells were quantified as described in Figure 5. n = 4-28 larvae/group.

**Figure 6- Supplemental Figure 1.**
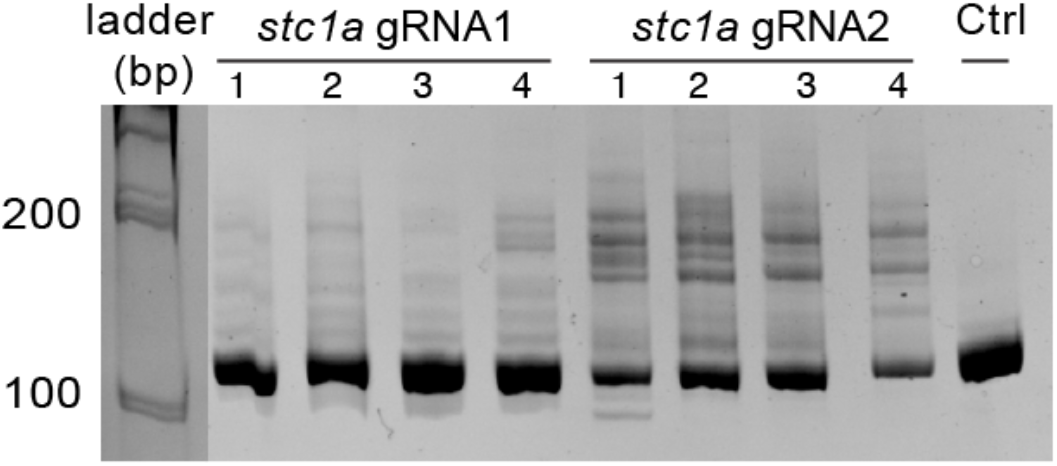
Validation of *stc1a* gRNA efficiency. Embryos injected with gRNAs and Cas9 mRNA were raised to 1 dpf. Each of them was lysed and analyzed by PCR followed by hetero-duplex motility assay.

**Figure 6- Supplemental Figure 2.**
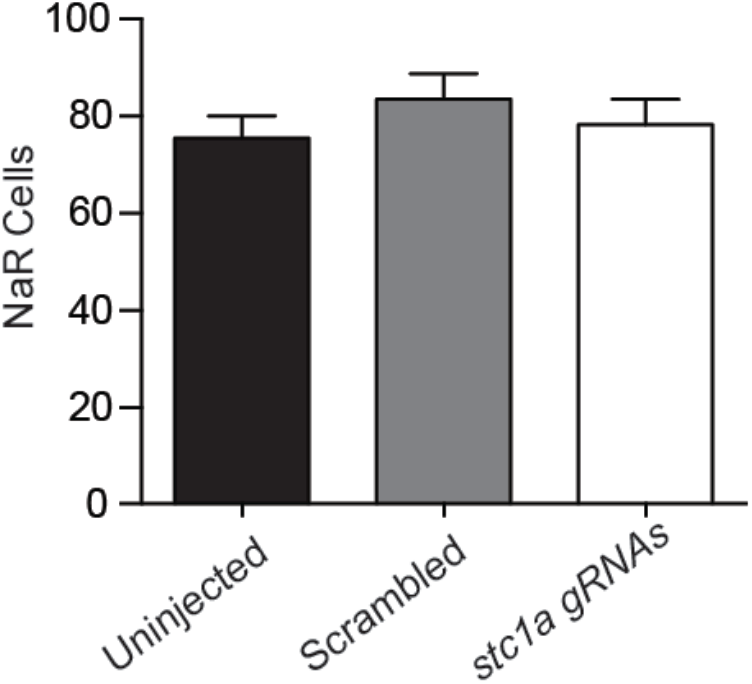
Igfbp5a is required in Stc1a regulation of NaR cell quiescence-proliferation balance. Progeny of igfbp5a^+/-^; *Tg (igfbp5a: GFP)* intercrosses were injected with scrambled gRNAs (Scrambled) or *stc1a* targeting gRNAs and Cas 9 mRNA. The injected embryos were raised in E3 embryo medium and analyzed at 5 dpf. n =39~43. No significant difference was detected among the groups.

